# A bottom-up approach for the *de novo* design of functional proteins

**DOI:** 10.1101/2020.03.11.988071

**Authors:** Che Yang, Fabian Sesterhenn, Jaume Bonet, Eva van Aalen, Leo Scheller, Luciano A Abriata, Johannes T Cramer, Xiaolin Wen, Stéphane Rosset, Sandrine Georgeon, Theodore Jardetzky, Thomas Krey, Martin Fussenegger, Maarten Merkx, Bruno E Correia

## Abstract

*De novo* protein design has enabled the creation of novel protein structures. To design novel functional proteins, state-of-the-art approaches use natural proteins or first design protein scaffolds that subsequently serve as templates for the transplantation of functional motifs. In these approaches, the templates are function-agnostic and motifs have been limited to those with regular secondary structure. Here, we present a bottom-up approach to build *de novo* proteins tailored to structurally complex functional motifs. We applied a bottom-up strategy to design scaffolds for four different binding motifs, including one bi-functionalized protein with two motifs. The *de novo* proteins were functional as biosensors to quantify epitope-specific antibody responses and as orthogonal ligands to activate a signaling pathway in engineered mammalian cells. Altogether, we present a versatile strategy for the bottom-up design of functional proteins, applicable to a wide range of functional protein design challenges.

---

*De novo* protein design has emerged as a powerful approach to expand the natural protein repertoire (*1–5*). The majority of previous studies have been focused on structural accuracy of computational models relative to experimentally determined structures and thermodynamic stability of the designs (*1–5*). In contrast, the design of *de novo* proteins encoding biochemical functions is lagging far behind (*6*). Nonetheless, successes to date illustrate the potential of *de novo* design to transform multiple areas of biology and biotechnology, including the design of vaccine candidates (*7–9*), protein-based lead drugs (*10*), antivirals (*11*), pH-responsive carriers (*12*) and others (*13–15*).

A widely used approach to design functional proteins is to transplant functional sites from their native context to heterologous proteins derived from the natural protein repertoire or *de novo* designed structures (*16–19*). Commonly, we refer to this two-step approach, consisting of first selecting or building a stable, functionless scaffold, which subsequently serves as template for grafting, as a ‘top-down’ approach (Fig. 1a).

**Fig. 1:**
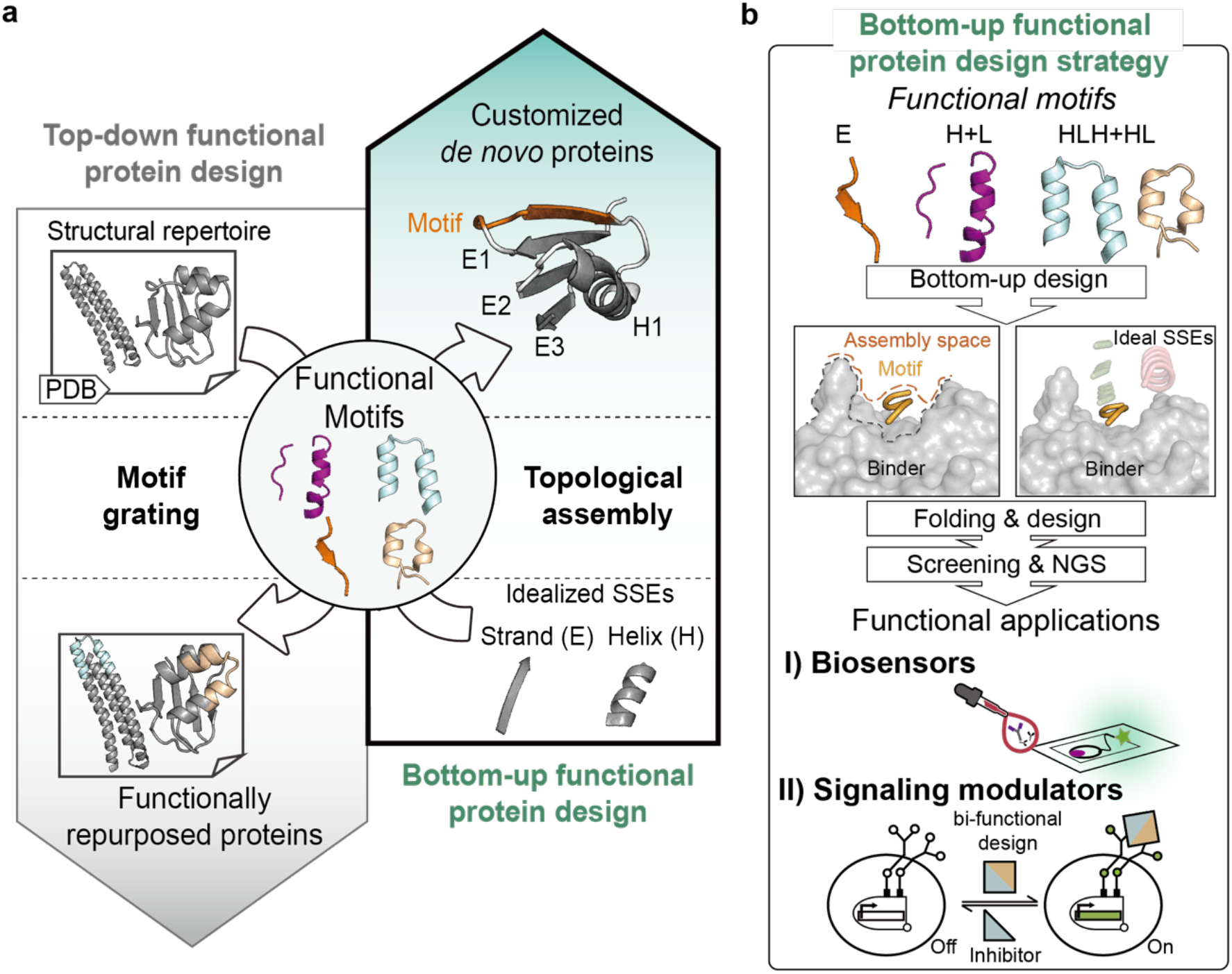
Bottom-up design of functional *de novo* proteins. **a**, Two conceptual frameworks for designing functional proteins. In top-down strategies, functional motifs are grafted onto structural templates found in the existent protein structural repertoire. In contrast, the presented bottom-up strategy assembles secondary structures around a given functional motif from scratch. **b**, The bottom-up design strategy was applied to design scaffolds for four different binding motifs, obeying spatial constraints imposed by the binding partner. To identify lead candidates for functional characterization, designed topologies were screened using yeast display, followed by next-generation sequencing (NGS). The designed proteins were functional as biosensors for the detection of epitope-specific antibodies in complex samples, and as signaling triggering molecules of synthetic receptors in engineered mammalian cells.

There are several important limitations of ‘top-down’ approaches for functional protein design. An essential prerequisite is the availability of template structures in the natural repertoire with enough local structural similarity to allow grafting of the functional site. Consequently, with a few exceptions (*20*), grafting has been largely limited to single, regular secondary structures that are frequently found in natural proteins (*11, 17, 19*). However, the vast majority of functional sites are not contained in single, regular helical segments, but rather are composed of multiple and often irregular structural segments that are stabilized by the overall protein structure (*21–23*).

Similarly, most *de novo* proteins are built with a high content of regular secondary structures, high contact order and minimal loops (*5*). While these proteins generally are thermodynamically very stable, *de novo* proteins designed in a ‘function-agnostic’ fashion are unlikely to have sufficient local structural similarity to an irregular, multi-segment, functional motif that would make them amenable to grafting approaches.

In contrast to this ‘top-down’ approach, a ‘bottom-up’ (Fig. 1a) strategy could consider the local structural and global topological requirements of the functional motif, irrespective of its complexity, and build supporting secondary structural elements (SSEs) to stabilize the functional site in its native conformation (*24*). A few studies have employed such a function-centric design strategy (*9, 10, 13, 25, 26*), but a general approach that allows the systematic construction of *de novo* proteins with embedded functional motifs, defined connectivity and precise control over the spatial positioning of each motif and secondary structure element is thus far lacking.

Here, we describe a ‘bottom-up’ approach to design functional *de novo* proteins. SSEs are assembled around the functional motifs of interest, followed by a folding-design stage where the motifs are kept static and the scaffold structure is further tailored for accurate presentation of the motif (*27*). Extending on our previous work (*9*), we demonstrate the power of this ‘function-centric’ approach by designing various protein folds (all-alpha, alpha with crossover, alpha-beta), to accommodate irregular and discontinuous antibody binding motifs derived from four viral epitopes. On the functional side, we showcase two completely distinct applications of *de novo* designed proteins (Fig. 1b). First, we show their utility as components of BRET-based biosensors to detect and quantify epitope-specific antibodies. Second, we assembled a protein topology with two distinct binding motifs, which was functional to trigger receptor-mediated signaling in engineered mammalian cells. Overall, we present a versatile strategy for the design of functional *de novo* proteins carrying structurally complex binding motifs, applicable to a wide range of functional protein design challenges.

## Results

### Bottom-up functional *de novo* protein design

We have recently explored the bottom-up conceptual approach, namely the TopoBuilder protocol, for the *de novo* design of epitope-focused immunogens (*9*). Here, we enlarge the protein structural space explored and the scope of the applications, by introducing a more generalizable technical framework for function-centric, bottom-up *de novo* design of functional proteins (Fig. 1).

The TopoBuilder enables the building of idealized protein structures tailored to a given functional motif (Fig. S1). Briefly, each topology is initially defined in a string format using the forms definition (*28, 29*) which are then translated into a two-dimensional topological representation of the protein. This compact protein representation scheme allows the rapid enumeration of different protein folds to stabilize a given functional motif. Subsequently, selected topologies are projected into the three-dimensional space, incorporating the coordinates of the functional motif to generate idealized 3D sketches. Secondary structures are placed with default distances of 10-11 Å between the center of mass of alpha-helices, and 4.5 – 5 Å between adjacent beta-strands. Pairwise distance constraints are derived from the idealized 3D sketches to guide folding simulations coupled to sequence design using Rosetta FunFolDes (*27*).

To showcase the capability of the proposed bottom-up strategy to build a variety of protein folds for structurally diverse functional motifs, we selected four well-characterized antibody binding motifs of the respiratory syncytial virus (RSV) F and G proteins, differing in secondary structure content and structural complexity (Figs. 1, 2): I) RSVF site IV, a linear, irregular beta strand (*30*); II) RSVF site 0, a discontinuous epitope consisting of a kinked alpha helix and a disordered loop (*31*); III) RSVF site II, a linear helix-turn-helix motif (*32*); IV) RSVG 2D10 epitope, a linear helix-loop segment that is constrained by two disulfide bonds (*33*).

**Fig. 2:**
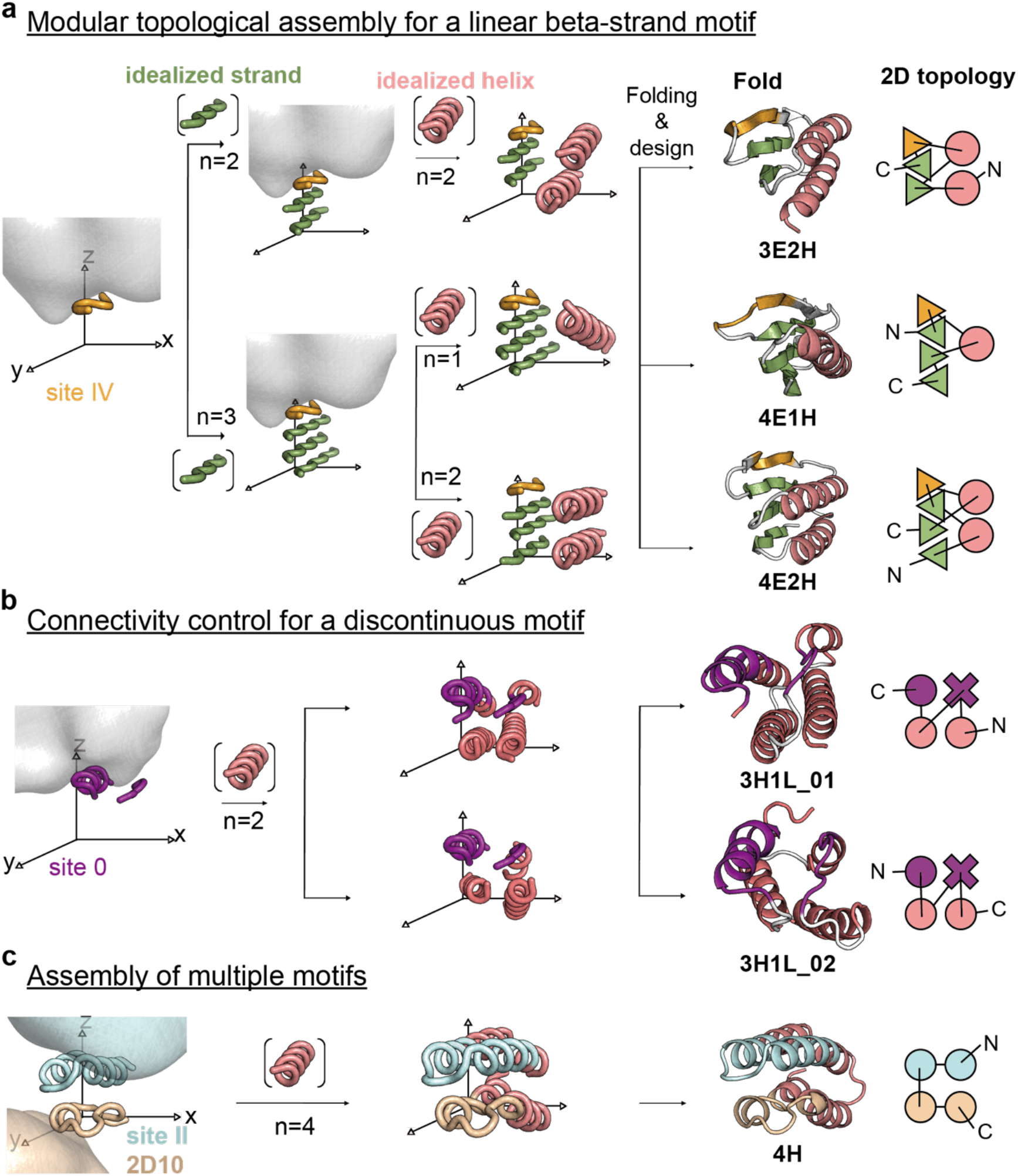
Bottom-up design of six protein folds for the presentation of four different binding motifs. **a**, Design of three different protein topologies tailored to the RSVF site IV epitope. The TopoBuilder enables a modular topological assembly of virtually any protein fold that can be described in layers, by controlling the spatial positioning of beta-strands and alpha-helices of defined length. **b**, The discontinuous RSVF site 0 was stabilized by two additional helices in two folds with different SSE connectivities. **c**, Building and design of a helical topology presenting two binding motifs (RSVF site II and RSVG 2D10 epitopes). Topologies are named according to their designed secondary structural content. Two dimensional topological representations indicating the connectivity are shown on the right for each fold. E: beta strand, H: alpha helix, L: loop structure.

As shown in Fig. 2, the TopoBuilder provides three major assets for functional protein design. First, it enables the modular assembly of de novo generated protein topologies with respect to the functional motif’s structural requirements. To do so, idealized SSEs with defined length are positioned to support the motif, enabling the systematic building and exploration of multiple topologies accommodating the same motif. Leveraging this feature, we constructed three different topologies to present the site IV epitope: 3E2H (three-stranded β-sheet and two helices), 4E1H (four-stranded β-sheet and one helix) and 4E2H (four-stranded β-sheet and two helices). Second, it allows full control over the topology’s connectivity, as exemplified by the design of two different connectivities for a 3H1L topology (three-helix bundle with a cross-over loop) assembled to host the discontinuous site 0 epitope. Third, the TopoBuilder enables the building of protein scaffolds accommodating multiple functional motifs. We assembled a four-helix bundle (4H topology) with the site II and the 2D10 epitope in a back-to-back orientation. Notably, searches on the Protein Data Bank (PDB) for design templates with local structural similarity to both site II and 2D10 motifs resulted in an extremely low number of potentially designable scaffolds (Fig. S2). This observation further highlights the value of the bottom-up approach for the design of functional proteins with structurally complex or multiple motifs.

Upon assembling the 3D sketch, typically 10,000 - 20,000 designs were generated for each topology with folding-design simulations (*27*). This initial pool of designs was filtered according to structural features such as well-packed hydrophobic cores, paired beta-strands, and sequences that presented funnel-shaped energy landscapes in Rosetta *ab initio* simulations (*34*). Based on the computational designs we constructed combinatorial sequence libraries as follows: surface residues were restricted to the amino-acid more frequently sampled in the designs; several boundary and core residues were also restricted to a single amino-acid based on the designs; remaining boundary and core residues were allowed limited variability encoding amino-acids observed in the computational designs (Figs. S3-S7). The combinatorial libraries yielded diversities of 10^6^-10^7^ sequences per topology.

Each library was sorted under double selective pressure: site IV or site 0 designs – antibody binding (101F and 5C4, respectively) and protease digestion to ensure that the designs were folded; bifunctional designs - binding to two different antibodies (Mota and 2D10) to ensure both motifs were presented in their native conformation. For each designed fold we sorted two populations, one under stringent conditions (i.e. best 1-2% after chymotrypsin treatment or low antibody labeling concentration) and another under permissive conditions (i.e. top 20% in the absence of protease treatment and high antibody labeling concentrations) (Fig. S8). These populations were bulk-sequenced using next-generation sequencing (Fig. S8) and enrichment scores for each sequence under stringent conditions were computed (see methods).

A comparison between sequences that showed positive enrichment scores for both stability and antibody binding versus those with negative enrichment scores revealed amino acid preferences for each sampled position in the different folds (Figs. S3 - S7). Upon modeling the 100 - 200 sequences with the strongest positive and negative enrichment scores, we found that three out of five topologies showed a higher side chain carbon-carbon contact number in the protein core, indicating that improved hydrophobic core packing was a critical determinant of successful designs (Fig. S9). However, this observation was not fully general as two of the topologies did not follow this trend (Fig. S9). When comparing the top scoring designed sequences from the initial pool to those that were experimentally enriched for stability and binding, the sequence identity of the sampled core positions was on average 47% (Table S1). Together, these results underline the challenge of designing functional *de novo* proteins purely based on computational predictions and highlights the concurrent need for experimentally sampling a diversified sequence space while selecting for function and stability. In contrast to several previous protein design efforts that employed saturation mutagenesis optimization of computational designs (*10, 11, 17*); here we designed focused libraries restricted to computationally predicted amino acids and those with similar chemical properties, which yielded well-folded and functional proteins in a single round of screening.

### *De novo* designs are well folded and bind with high-affinity to target antibodies

For each topology, 5-12 designed sequences that were strongly enriched under double selection pressures in the yeast screen were selected for recombinant expression in *E.coli* and subsequent biophysical characterization.

All three topologies designed to accommodate site IV yielded well-folded and stable proteins that bound the 101F antibody with dissociation constants (K_D_) ranging from 20 to 180 nM (Fig. 3 and Figs. S10 - S12). While these affinities are slightly lower than that of the 101F-RSVF interaction (K_D_ = 4 nM), they are substantially higher compared to the peptide epitope (KD ~ 60 μM (*30*)), indicating the epitope conformation is accurately presented. The best designs from each design series, 3E2H.37, 4E1H.95 and 4E2H.210, showed melting temperatures (T_m_s) of 74 °C, 58 °C and 84 °C, respectively (Fig. 3).

**Fig. 3:**
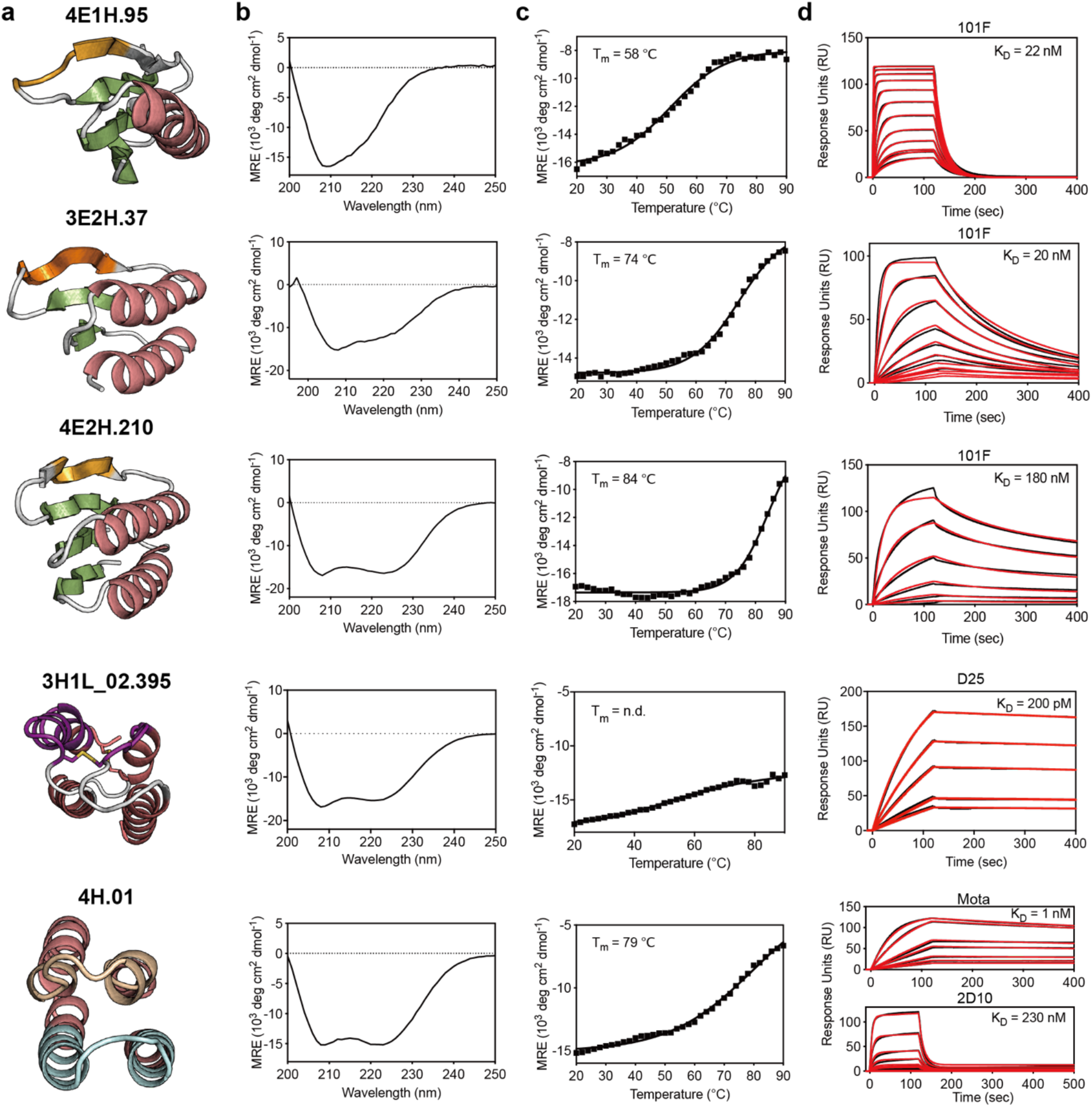
Biophysical characterization of lead variants from each topology. **a**, Computational models of lead designs for each topology. **b**, Designs are well-folded and show circular dichroism (CD) spectra in agreement with the secondary structure content of the designed proteins. **c**, Designed proteins are thermally stable as measured by CD. **d**, Binding affinity to target antibodies determined by SPR. Sensorgrams are shown in black, fitted curves in red. Binding of 4E1H.95 and 4E2H.210 was measured against immobilized 101F IgG, 3H1L_02.395 against D25 IgG, and 4H.01 against both Motavizumab (Mota) and 2D10 IgG. n.d., non-determined.

In terms of connectivity exploration within the TopoBuilder, both variants of the 3H1L topology, presenting the discontinuous site 0 epitope, were recombinantly expressed and bound to their target antibody D25. However, all sequences tested from the 3H1L_01 design series were dimeric in solution, and failed to bind to the 5C4 antibody (Fig. S13). The 5C4 antibody was shown to engage site 0 from a different angle compared to D25 (*35*), and binding to both antibodies can thus serve as a probe for the conformational integrity of the discontinuous site 0. In contrast, the 3H1L_02 designs bound both D25 and 5C4, with affinities of 0.2 - 0.9 nM for D25 and 7 - 25 nM to 5C4 (Fig S14). The best design of the 3H1L_02 series (3H1L_02.395) bound to D25 and 5C4 with K_D_s of 0.2 nM and 25 nM, respectively (Fig. 3 and Fig. S15). Both binding affinities closely match the reference binding affinity of the respective antibodies binding to prefusion RSVF (^*D25*^K_D_= 150 pM; ^*5C4*^K_D_ = 13 nM), indicating that this discontinuous motif was mimicked accurately in *de novo* designed proteins (*31*). Also, 3H1L_02.395 was monomeric in solution, thermostable with only partial unfolding at 90 °C, and showed a well-dispersed HSQC NMR spectrum indicating that the protein is well folded (Fig. 3 and Fig. S15).

For the 4H topology carrying two functional motifs, 8 out of 12 designs were soluble upon purification, showed CD spectra typical of alpha-helical proteins and bound to both target antibodies, Motavizumab (Mota) and 2D10, with K_D_s ranging from 0.5 to 10 nM for Mota and 200 to 400 nM for 2D10 (Fig. S16). The best variant, 4H.01, was well folded and thermostable according to CD and NMR, showing a T_m_ of 75 °C (Fig. 3, Fig. S16). 4H.01 bound with a K_D_ of 0.5 nM to Mota and 200 nM to 2D10 (reference affinities to native antigens ^*Mota*^K_D_= 30 pM; ^*2D10*^K_D_ = 10 nM). Overall, these results confirm that the presented ‘bottom-up’ strategy can design *de novo* proteins carrying complex functional motifs accurately presented in stable protein folds.

### Experimentally structures confirm the accuracy of the computational designs

Next, we solved a crystal structure of 4E1H.95 in complex with the target antibody (101F) at 2.9 Å resolution. The structure is in close agreement with the computationally designed model, with a backbone RMSD of 2.5 Å (Fig. 4a). The designed backbone hydrogen bonding network and the overall pairing of the β-sheet was accurately recapitulated in the crystal structure as compared to the computational model (Fig. 4a). In the bound state, the β-bulged structure of the site IV epitope also retained the designed hydrogen bond network supported by an adjacent beta strand. Upon superposition to the antibody-peptide bound structure (*30*), 4E1H.95 mimicked the site IV with sub-angstrom accuracy (Fig. 4b).

**Fig. 4:**
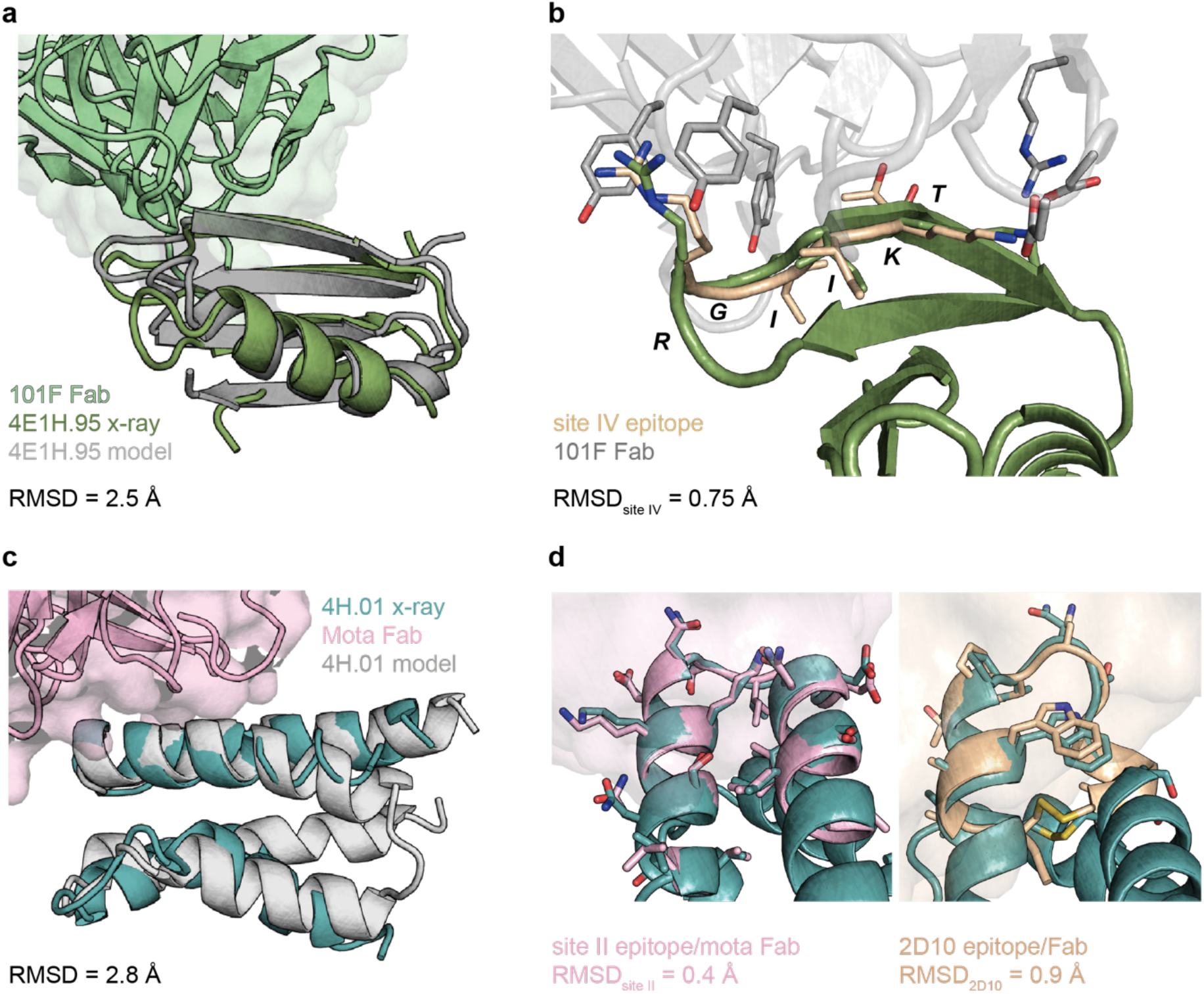
Crystal structures are in close agreement with the design models. **a**, The crystal structure of 4E1H.95 in complex with the 101F Fab (green surface) was solved at 2.9 Å resolution, and closely resembles the design model (RMSD_backbone_ = 2.5 Å). **b**, Superposition of 4E1H.95 crystal structure with the antibody-bound site IV peptide epitope (PDB 3O41) shows accurate mimicry of the designed conformation (RSMD_backbone_ = 0.9 Å). **c**, The crystal structure of the bifunctional 4H.01 scaffold in complex with mota Fab (pink surface) aligns to the computational model with a backbone RMSD of 2.8 Å. **d**, The 4H.01 structure closely matches the site II epitope (pink, PDB 3IXT, RMSD_backbone_ = 0.4 Å) as well as the 2D10 epitope (wheat, PDB 5WN9, RMSD_backbone_ = 0.9 Å).

We also solved a crystal structure of 4H.01 in complex with the Mota Fab at 2.8 Å resolution. The 4H.01 scaffold closely resembles the design model, with a backbone RMSD of 2.8 Å (Fig. 4c). The site II epitope, in its bound form, accurately matches the epitope with a backbone RMSD of 0.4 Å (Fig. 4d). Similarly, the 2D10 epitope in 4H.01 shows a full-atom RMSD of 0.9 Å to the 2D10 epitope of RSVG (Fig. 4d), providing a rational basis for the high-affinity binding observed. Together, these structures confirm that the TopoBuilder design pipeline yields proteins that closely resemble the computational models while preserving the structural integrity of several functional motifs.

### Biosensors containing *de novo* designs detect epitope-specific antibodies

In recent years, important studies in the vaccine field have revealed correlations between antibody specificity and the degree of protection afforded against several pathogens upon vaccination or natural infection (*36, 37*). However, standard assays mostly measure bulk serum titers, failing to provide a detailed picture about the epitope specificities. The detection and quantification of epitope-specific antibodies is currently performed by competition assays with monoclonal antibodies, which are laborious and often require extensive empirical optimization (*38*). In conjunction with the fast-growing collection of atomic-level structures of neutralizing antibodies and their target epitopes to inform the design of precision immunogens (*39–41*), assays to dissect epitope-specific responses in bulk sera may help to improve our understanding of natural and vaccine-induced immunity.

We used the neutralizing epitopes identified in RSVF as a test case and hypothesized that a biosensor based on the designed scaffolds would be an attractive tool to detect and quantify epitope-specific antibodies in bulk serum. We designed a biosensor based on the recently developed LUMABS platform, as shown in Fig. 5a (*42*). Briefly, the sensor is based on bioluminescence resonance energy transfer (BRET) where in its “closed” conformation, BRET occurs between the nanoluciferase (Nluc) donor and the mNeonGreen (mNG) acceptor. In the presence of antibodies specific for the presented epitope, the sensor adopts an open state, as indicated by a decrease in the BRET ratio (Fig. 5b).

**Fig. 5:**
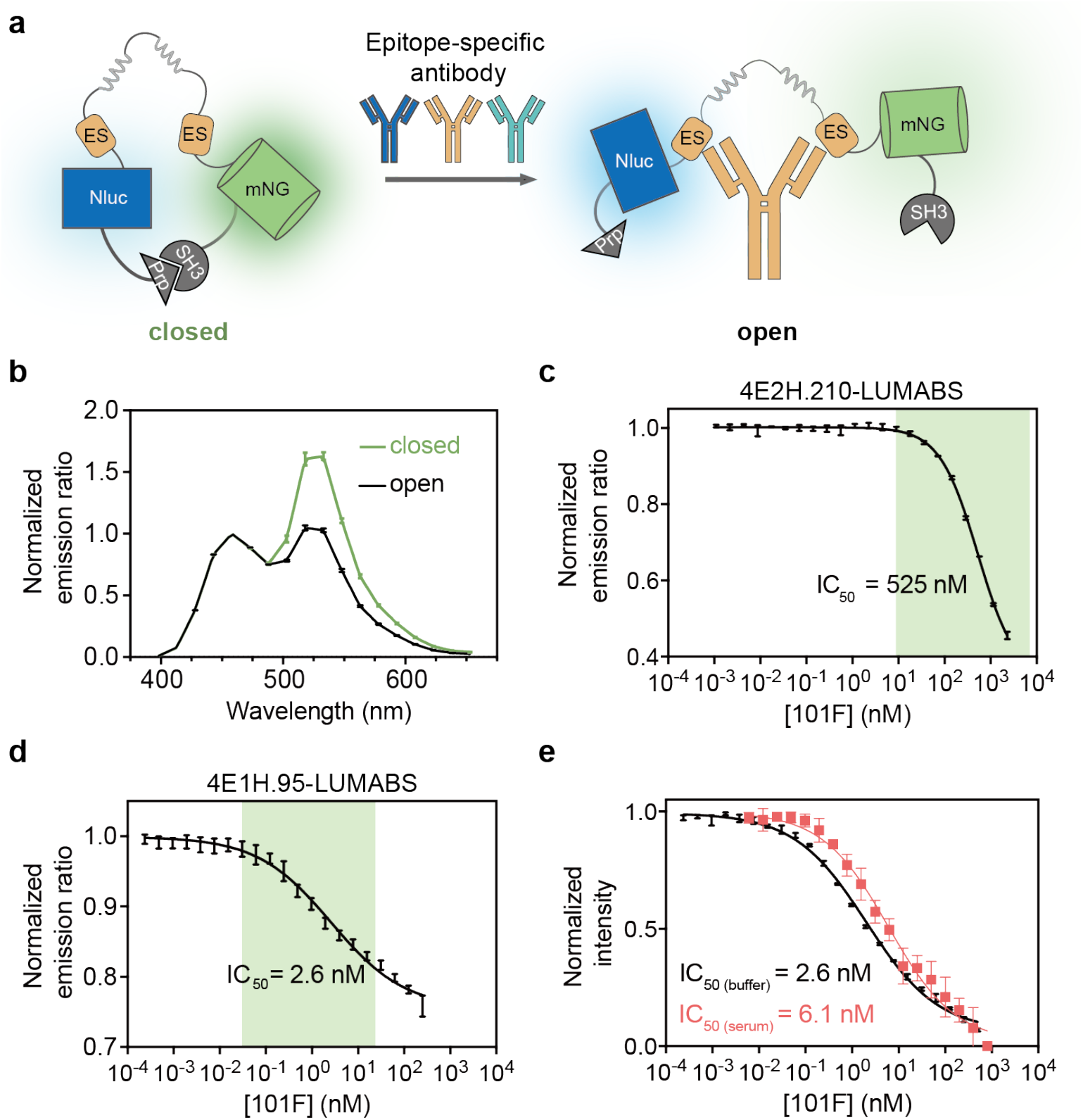
Antibody biosensors based on *de novo* designed proteins for the detection of site-specific responses. **a**, Schematic representation of the LUMABS sensor platform. NanoLuc luciferase (Nluc) and mNeonGreen (mNG) are held in close proximity by two helper domains (SH3 domain and a proline-rich-peptide, Prp), allowing efficient BRET between Nluc and mNG. Antibodies binding to the designed epitope-scaffolds (ES) disrupt the weak Prp-SH3 interaction, opening the sensor and decreasing BRET efficiency. The fluorescence signals were normalized with the emission intensity of Nluc. **b**, Luminescence spectra of the sensor in a closed (green) and open conformation. **c**, Characterization of site IV sensor based on the 4E2H.210 scaffold upon titration of 101F IgG. The plot shows the ratio between 518 and 458 nm emission upon titration of 101F antibody, yielding an IC_50_ of 525 nM (green: detection range). **d**, Characterization of site IV sensor based on the 4E1H.95 scaffold upon titration of 101F IgG. In agreement with the higher affinity compared to 4E2H.210, 4E1H.95 showed an IC_50_ of 2.6 nM. **e**, The 4E1H.95_LUMABS shows similar performance in human plasma (red) compared to buffer. Data points are represented as mean ± SEM.

As a proof-of-concept, we developed sensors for the RSVF antigenic site IV, an important neutralization epitope that is conserved between two major respiratory pathogens, RSV and metapneumovirus (MPV), and thus targeted by virus cross-neutralizing antibodies (*43*). To do so, we embedded either the 4E1H.95 or the 4E2H.210 designs in the LUMABS sensor, which differ in affinity for the 101F antibody by approximately one order of magnitude. The sensors were robustly expressed in *E.coli*, and showed a concentration-dependent decrease in BRET signal upon the addition of the 101F antibody (Fig. 5c,d), but not in response to a control IgG (Fig. S18). In agreement with the different affinities for the target antibody, the 4E1H.95-based LUMABS was able to detect 101F antibody present at sub-nanomolar concentrations (<0.1 μg/ml) up to ~50 nM, whereas 4E2H.210 showed a dynamic range between ~30 nM up to micromolar concentrations (Fig. 5c,d). Thus, by constructing LUMABS sensors based on scaffolds with different affinities, the epitope-specific antibody quantification can be extended over a large range of antibody concentrations. Importantly, when tested directly in human sera spiked with 101F antibody, a complex sample with high concentrations of immunoglobulins with diverse specificities, the sensor was functional and showed very similar performance compared to its performance in buffer (Fig. 5e).

Together, the presented sensors could be versatile tools for immune monitoring, enabling the study of serum responses beyond bulk titers, and the quantification of the levels of epitope-specific antibodies elicited by viral infection or vaccination.

### Bi-functionalized *de novo* designs regulate the activity of synthetic cell-surface receptors

Many fundamental biological processes at the cellular and organismal levels are regulated by the interaction of soluble effector proteins with cell surface receptors. Within the realm of synthetic biology, substantial interest has emerged in rationally engineering receptor-ligand pairs to control cellular behavior (*44–46*). Given that engineered receptors which can be triggered by *de novo* proteins should be orthogonal to endogenous ligand-receptor pairs, we sought to create a synthetic ligand-receptor system that modulates the heterodimerization of synthetic receptors using our bi-functional *de novo* design (4H.01) (Fig. 6a).

**Fig. 6:**
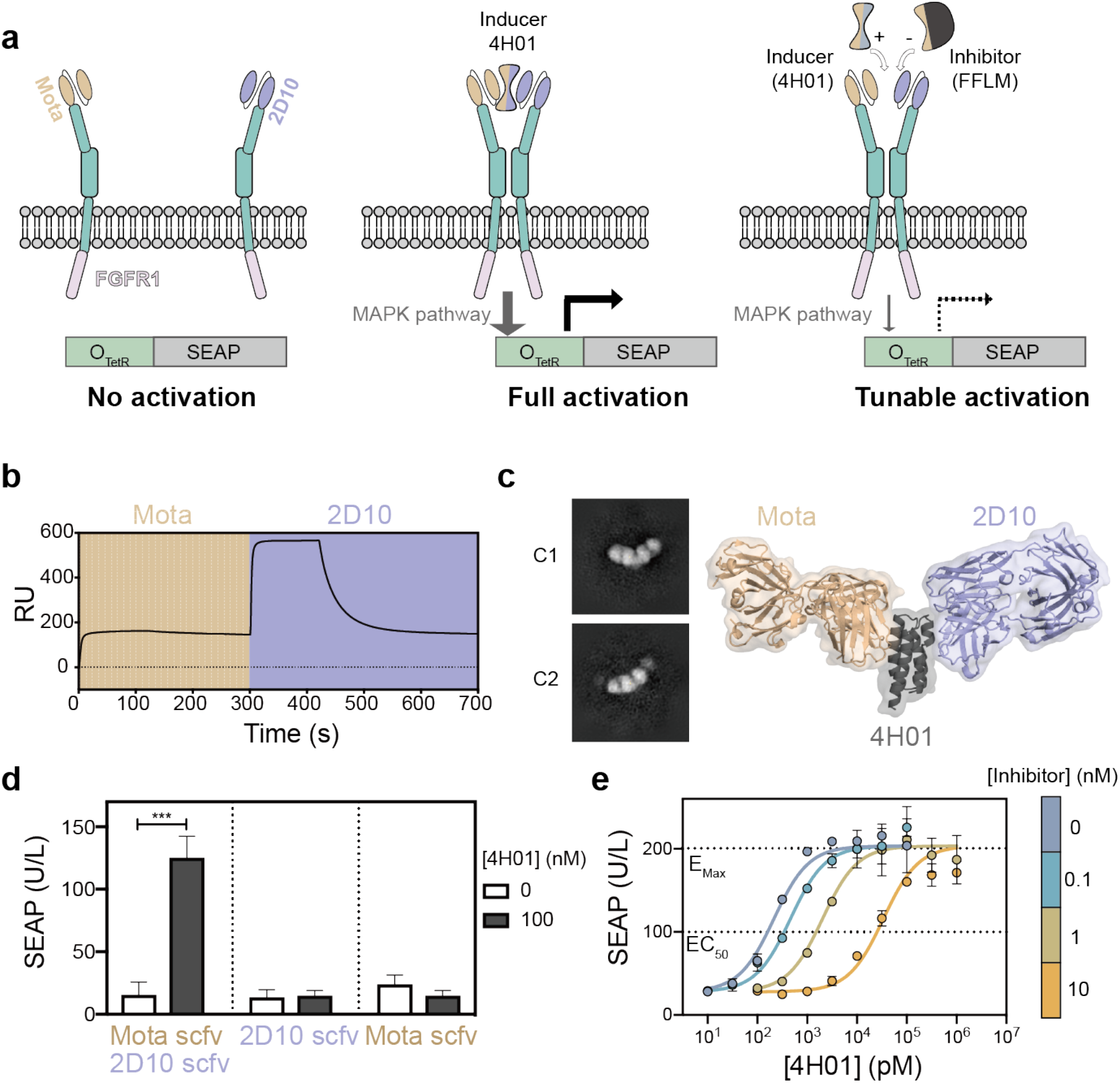
Bi-functional *de novo* design controls the activity of synthetic receptors in mammalian cells. **a**, Schematic representation of the receptor architecture and the regulation of the signaling activity monitored through the expression of a reporter gene (SEAP). Two scFvs were used as ectodomains of the synthetic receptors which hetero-dimerize in the presence of 4H.01, resulting in activation of the MAPK signaling pathway that controls the expression of the reporter gene. In the presence of a computationally designed high-affinity antagonist carrying only the site II epitope, the heterodimerization is prevented. **b**, SPR binding response of 4H.01 to target antibodies. 4H.01 binds simultaneously to Mota and 2D10, as shown by the 2D10 binding signal in presence of saturating amounts of Mota. **c**, EM images of two 2D average classes of the 4H.01 design complexed with both mAbs (left), supporting that both antibodies can bind simultaneously as suggested by the computational model (right). **d**, Responsiveness of dual and single receptor expressing cells. 15 hours after transfection, cells were treated with 100 nM of 4H.01. The dose-response rate was quantified by SEAP expression after 24 hours. **e**, Inhibitory effect of a computationally designed antagonist. Transfected cells were treated with different molar ratios of 4H.01 and FFLM for 24 hours and SEAP production was quantified. The EC_50_ was determined by fitting the SEAP response with the dose-response equation implemented in GraphPad Prism 8.0 (goodness of fit greater than R^2^ > 0.95). Data shown in d) and e) were obtained in three independent replicates. *** p = 0.0007, unpaired Student’s t-test.

We used the recently developed synthetic receptor platform developed by Scheller and colleagues (*47*), on which we fused scFvs (single-chain variable fragments) of Mota and 2D10 mAbs to the extracellular domains of EpoR (erythropoietin receptor). Intracellularly, the EpoR scaffold contained the FGFR1 (fibroblast growth factor receptor 1), leading to the activation of the endogenous MAPK pathway and expression of the reporter protein SEAP (human placental secreted alkaline phosphatase). In this system, ligand-induced heterodimerization of the Mota/2D10-EpoR receptors can be directly quantified by measuring SEAP activity (Fig. 6a).

As predicted by the structural model of the design, we confirmed that 4H.01 can bind simultaneously to both mAbs by SPR and by negative stain electron microscopy (Fig. 6b-c). We then introduced the synthetic receptors (scfv_Mota_-EpoR-FGFR1 and scfv_2D10_-EpoR-FGFR1) into HEK cells and measured SEAP expression upon titration of 4H.01. Signaling activation was observed when the engineered cells expressed both extracellular scFv-fused receptors and the response curve revealed a typical dose-dependent activation with a high sensitivity (EC_50_ = 214 pM; 95% CI = 166 – 276 pM) (Fig. 6d).

Cellular signaling is often the result of a delicate balance between agonists and antagonists. To tune the signaling output induced by our *de novo* designed protein, we used a previously designed protein (FFLM) (*8*) presenting only the site II epitope to function as an antagonist. FFLM binds to Mota with approximately one order of magnitude higher affinity compared to 4H.01 (K_D_ ~ 20 pM) (*8*). We tested the response of the synthetic receptors to mixtures of designed agonists and antagonists used at different molar ratios (Fig. 6a), and observed an antagonist dose-dependent response with increasing amounts of the inhibitor FFLM (Fig. 6e). The effective EC_50_ was shifted correspondingly from 214 pM (in the absence of inhibitor) to 459 pM ([inhibitor] = 100 pM), 2 nM ([inhibitor] = 1 nM) and 34 nM ([inhibitor] = 10 nM). The ability of fine-tuning receptor activity by the action of computationally designed agonist and antagonist may unveil new options for encoding complex regulatory mechanisms (e.g. feedback loops) for precision-guided cell-based therapeutics.

Altogether, we have demonstrated that the presented bottom-up design approach is well suited to design functional proteins with irregular, discontinuous as well as multiple binding sites, which is inherently difficult using top-down approaches.

### Discussion

The design of novel functional proteins is a fundamental test to our understanding of protein structure and function, and has enormous potential in multiple arenas in basic biology and biotechnology (*7–15, 17*). Nevertheless, the majority of studies where function was the main objective, existing proteins were repurposed and used as templates to install functional sites (*7, 17–20*). This ‘top-down’ approach faces inherent limitations for functional motifs with high structural complexity (e.g. irregular conformations or multi-segment motifs), as the number of designable templates available in the natural protein repertoire becomes sparse. To overcome this critical bottleneck, we present a ‘bottom-up’ approach for the *de novo* design of functional proteins.

In our motif-centric *de novo* design approach, we account for the structural and sequence constraints of the functional motifs, serving as a ‘functional seed’ to build tailored scaffolds that stabilize their native conformation. While previous reports have described conceptually similar ‘bottom-up’ design approaches of functional proteins, only one topology per functional site was reported, and the topologies were structurally limited to helical bundles (*10, 13, 25, 26, 48*).

The bottom-up strategy presented here has significant advantages for the design of *de novo* functional proteins. We have successfully designed different *de novo* topologies presenting a diverse set of structural motifs, including those that are irregular and include multiple segments. We attempted the design of six different topologies, five of which yielded well-folded and stable proteins. The functional sites were presented accurately in the *de novo* proteins, as shown by the high affinity binding to their target antibodies, as well as by two experimentally determined structures that confirmed the accuracy of the design models at the atomic level.

On the functional significance of our work, one of the most attractive prospects for *de novo* design is to create proteins with activities that are out of reach for natural proteins. Herein, we have shown the utility of the TopoBuilder as a motif-centric design strategy to accomplish this task.

First, the designed RSV epitope-presenting proteins enabled the engineering of functional biosensors, to detect and quantify epitope-specific antibodies in complex samples. Monitoring epitope-specific antibody responses at the serum level is of utmost importance to decipher immune responses upon natural infection, and to assess the efficacy of novel vaccine candidates (*41, 49*). Thus, the presented biosensors may become valuable diagnostic tools to dissect serum responses beyond bulk serum titers.

Second, we have shown the utility of the TopoBuilder for the *de novo* design of proteins presenting multiple binding sites, a common feature of natural proteins but largely unachieved in the field of *de novo* protein design. We have shown the potential of a bi-functional *de novo* protein to mediate the heterodimerization of synthetic receptors, resulting in the activation of a signaling pathway in synthetic cells. We foresee that the ability of designing receptor ligands with precise spatial configurations of the binding sites will expand the repertoire of orthogonal ligands for engineered cells.

Altogether, the presented bottom-up functional design strategy is an important step forward on the design of functional proteins. While herein the functions encoded were limited to molecular recognition mediated by antibody-binding motifs, the approach is widely applicable to other known functional motifs, and ultimately, extendable to the design of novel functions absent in natural proteins.

## Contributions

CY, FS and BEC conceived the work and designed the experiments. CY and FS performed computational design and experimental characterization. JB developed the TopoBuilder. EvA performed work related to biosensors, and LS performed cellular assays. CY solved x-ray structures and LAA performed NMR characterization. JTC, XW, SR, and SG performed experiments and analyced data. FS, CY and BEC wrote the manuscript, with input from all authors.

## Data availability

All code used for this study is available through a public Github repository: https://github.com/lpdi-epfl/Bottom-up-de-novo-design. Structures have been deposited in the Protein Data Bank.

## Acknowledgements

We thank Kelvin Lau, Aline E. Christine and Florence Pojer in PTPSP facility at EPFL for crystallography support. We thank the protein expression core facility (David Hacker, Laurence Durrer, Soraya Quinche) for help with mammalian protein expression, as well as Davide Demurtas from CIME and Sergey Nazarov from PTBIOEM (EPFL, Lausanne, Switzerland) for electron microscopy support. We thank the flow cytometry core facility at the EPFL for technical support and the gene expression core facility for help with next-generation sequencing. The computational simulations were facilitated by the CSCS Swiss National Supercomputing Centre as well by SCITAS at EPFL.

## Materials and Methods

### Computational design

The computational design procedure consisted of three main stages described below, which include several steps of backbone building, sequence design and selection.

#### Stage I) Topological sampling and backbone generation

We previously described the TopoBuilder (*9*) a python library for the construction of tailored protein scaffolds around structural motifs of interest. The TopoBuilder protocol is composed of three main steps (Fig. S1):

a. 2D definition of protein topologies – 2D topologies that are compatible with the motif of interest are defined using the forms scheme previously reported by Taylor and colleagues (*28*). The forms allow for the definition of multi-layer protein topologies using string-based descriptors that define the secondary structure elements (SSEs) and the connectivity between them.
b. 3D projection of selected topologies – 2D forms compatible with topological and functional requirements (e.g. connectivity, chain directionality, no crossover loops) of the motifs are projected in 3D. To do so, idealized SSEs of user-defined lengths are assembled in a 3D coordinate system, with user-defined distances and relative spatial orientations between all SSEs. Default distance between alpha-helices was 10 Å, and adjacent beta-strands was 4.5 Å. The resulting 3D structures are referred to as sketches.
c. Diversification of sketches using parametric sampling – the TopoBuilder uses parametric sampling to apply rigid body transformations to the individual SSEs. By doing so, variants of the sketches that maintain the same topology but have distinct orientations and distances between the SSEs can be rapidly sampled.

#### Stage II) Full-atom modeling and sequence design

To refine the sketches at the all-atom level, we use Rosetta FunFolDes as described previously (*27*). Briefly, the sketches served as templates to guide simulations performing constrained folding coupled with sequence design. In these simulations the conformation of the functional motif is kept static while the overall protein is folded and adapted to fit the structural constraints of the motif. Layer-based FastDesign (*50*) was used to design protein sequences. Each position was assigned a layer (core, surface or boundary) and a secondary structure type (helix, loop or beta strand) to restrict the type of amino acid sampled during design. For each designed topology between 10,000 and 20,000 designs were generated.

#### Stage III) Design optimization and selection

Designs were filtered according to multiple features, such as core packing, RMSD drift of the epitope after relax simulations and the match of secondary structure prediction to the design topology. The design filtering steps were performed using the rstoolbox python library (*34*). A subset of sequences was evaluated according to their propensity to recover the structure of the designed fold in Rosetta *ab initio* simulations. In some of the designed topologies, connecting loops between secondary structures were shortened using Rosetta Remodel (*51*) and disulfides were engineered using Rosetta Disulfidize (*52*). Command lines and RosettaScripts for the designed simulations are available in a Github repository (https://github.com/LPDI-EPFL/Bottom-up-de-novo-design/).

### Sequence selection for combinatorial screening using yeast surface display

We evaluated sequences generated by Rosetta, and selected 8-14 critical hydrophobic core positions and surface positions close to the binding interface in each designed topology to construct a combinatorial sequence library for high-throughput experimental screening (Fig. S3-S7). Combinatorial sequence libraries were assembled by PCR using DNA oligos (Table S3) containing degenerate codons at selected positions for combinatorial sampling of a selected subset of amino acids sampled by Rosetta, which were in some cases, diversified by additional amino acids with similar chemical properties. The theoretical diversity of the libraries ranged from 1×10^6^ to 1×10^7^. DNA oligos were diluted to a concentration of 10 μM, and assembled in a PCR reaction (55 °C annealing for 30 sec, 72 °C extension time for 1 min, 25 cycles). The full-length product was amplified in a second PCR. The PCR product was desalted and transformed into EBY-100 yeast strain as described previously (*53*), yielding a transformation efficiency of at least 1×10^7^ transformants. The transformed cells were grown for three days in SDCAA medium (1.25 g/L yeast nitrogen base, 5 g/L casamino acids, 13.6 g/L Na_2_HPO_4_, 8.5 g/L NaH_2_PO_4_, and 20 g/L glucose in 1 liter of deionized water) before induction. To induce cell surface expression, cells were pelleted, washed and resuspended in 100 ml SGCAA medium (SDCAA with galactose instead of glucose) with a cell density of 1 × 10^7^ cells/ml. After overnight induction, cells were washed three times in cold wash buffer (PBS + 0.05% BSA) and labelled with target antibodies 101F, D25, 5C4, Mota and 2D10 for 1 hour at 4 °C. Cells were washed twice with wash buffer and then incubated with FITC-conjugated anti-cMyc antibody and PE-conjugated anti-human Fc (BioLegend, #342303) or PE-conjugated anti-Fab (Thermo Scientific, #MA1-10377) for additional 30 min. Sorting was performed using a SONY SH800 flow cytometer in ‘ultra-purity’ mode. The sorted cells were recovered in SDCAA medium, and grown for 1-2 days at 30 °C.

To screen for protease resistant sequences as a proxy for well folded proteins, induced cells were washed three times with TBS buffer (20 mM Tris, 100 mM NaCl, pH 8.0) and resuspended in 0.5 ml of TBS buffer containing 0.5 - 1 μM of chymotrypsin. After five minutes at 30 °C, the reaction was quenched by adding 1 ml of wash buffer, followed by three to five wash steps. Cells were subsequently labelled with primary and secondary antibodies as described above.

### Next generation sequencing (NGS)

After sorting in different selection conditions, yeast cells were cultured overnight at 30 °C. Around 50 ml of yeast culture was harvested and DNA was extracted using Zymoprep Yeast Plasmid Miniprep II (Zymo Research). Two rounds of PCR were performed to amplify the coding sequence of the selected proteins and attach standard Illumina sequencing adaptors. The final amplified fragments were desalted (Qiaquick PCR purification kit, Qiagen) and analyzed for purity on a 2% agarose gel. Each library was sequenced using an Illumina NextSeq 2×150 bp paired end sequencing protocol (300 cycles), resulting in 1-2 million reads/sample.

For the data analysis, raw reads were trimmed and translated to the correct reading frame. For each sequence, an enrichment score was computed, based on its frequency under high-stringency selection versus low-stringency selection:

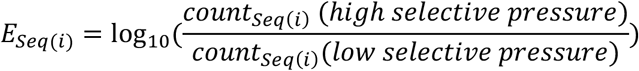

Positive enrichment values indicate a greater frequency of a given sequence under stringent selection conditions versus a low selective pressure. Between 10 and 20 sequences that were strongly enriched under two different selection pressures, such as high-affinity binding to two different antibodies or, alternatively, high-affinity binding to one antibody and chymotrypsin resistance, were recombinantly expressed and biophysically characterized.

### Protein expression and purification

#### Designed proteins

DNA was purchased from Twist Bioscience as DNA fragments, which were cloned into pET11b or pET21b expression vectors using Gibson cloning. A 6x His tag was added at the C terminus of 4E1H, 4E2H, 3E2H, 4H and 3H1L_02-based designs, and at the N terminus of the 3H1L_01 design series. Plasmids were transformed into *E.coli* BL21 (DE3) (Merck), and grown overnight in LB media supplemented with 100 μg/ml ampicillin. Overnight cultures were diluted 1:50 in TB medium and grown at 37 °C until the OD_600_ reached 0.6-0.8. To induce expression, 1 mM of isopropyl β-d-1-thiogalactopyranoside (IPTG) was added and cells were grown for 12-16 hours at 22 °C. Cultures were harvested and resuspended in lysis buffer (50 mM Tris, pH 7.5, 500 mM NaCl, 5% glycerol, 1 mg/ml lysozyme, 1 mM PMSF, 1 μg/ml DNase), and lysed by sonication. The cell lysate was pelleted by centrifugation (20,000 rpm, 20 mins) and supernatant was filtered with a 0.22 μm filter before loading onto a 1 ml HisTrap HP column (GE Healthcare). Proteins bound to the column were washed with 10 column volumes of washing buffer (50 mM Tris, pH 7.5, 500 mM NaCl, 10 mM imidazole) and eluted in 5 column volumes of elution buffer (50 mM Tris, pH 7.5, 500 mM NaCl, 300 mM imidazole). Eluted proteins were further purified by size exclusion chromatography on a Hiload 16/600 Superdex 75 pg column (GE Healthcare) in PBS buffer.

#### Antibodies and Fab fragments

For antibodies expressed as IgGs, the heavy and light chains encoding the variable domains were cloned into pFUSEss-CHIg-hG1 and pFUSEss-CLIg vectors (Invivogen), respectively. The antigen-binding fragments (Fab) were cloned into pHLSec vector (Addgene, #99845) using AgeI and XhoI restriction sites. Expression plasmids for heavy and light chains were mixed in a 1:1 stochiometric ratio before transient transfection of HEK293 cells as described previously (*8*). The collected supernatant was purified using a 1 ml HiTrap Protein A HP column (GE Healthcare) for IgG expression and 5 ml kappa-select column (GE Healthcare) for Fab purification. Bound antibodies/Fabs were eluted with 0.1 M glycine buffer (pH 2.7), immediately neutralized by 1 M Tris ethylamine buffer (pH 9), and buffer exchanged to PBS.

### SPR measurements

SPR measurements were performed on a Biacore 8K (GE Healthcare) at room temperature with HBS-EP+ running buffer (10 mM HEPES pH 7.4, 150 mM NaCl, 3 mM EDTA, 0.005% v/v Surfactant P20, GE Healthcare). IgGs were immobilized on CM5 chips (GE Healthcare # 29104988) via amine coupling (approximately 1000 - 1500 response units). The designed monomeric proteins were injected as analyte in two-fold serial dilutions at a flow rate of 30 μl/min, with a contact time of 120 seconds followed by 400 seconds dissociation time. After each injection cycle, surface regeneration was performed with 3 M magnesium chloride (101F ligand) or 0.1 M Glycine at pH 2.0 (Motavizumab, 2D10 and D25 ligands). Data were fitted with 1:1 Langmuir binding kinetic model using the Biacore 8K analysis software (GE Healthcare).

### Size-exclusion chromatography coupled with multi-angle light scattering (SEC-MALS)

SEC-MALS was performed on a HPLC system (Thermo Fisher) connected to a light scattering detector (miniDAWN TREOS, Wyatt). 100 μl of freshly purified protein (concentration 1-2 mg/ml) was injected on a Superdex 75 300/10 GL column (GE Healthcare) at a flow rate of 0.5 ml/min. UV absorption and light scattering were recorded and processed using the ASTRA software (version 6.1, Wyatt).

### Circular Dichroism

All circular dichroism data were collected on a Chirascan CD spectrometer (Applied Photophysics) using a quartz cuvette with path length of 1 mm. Purified proteins were diluted in 10 mM sodium phosphate buffer pH 7.4 to a final concentration of 30 μM. Far UV spectra were recorded between a wavelength of 190 nm and 250 nm with a scanning speed of 20 nm/min. The spectra were averaged from two repeated measurements and corrected for buffer absorption. To determine the thermostability of the designed proteins, temperature was ramped stepwise from 25 °C to 95 °C in increments of 2 °C in the presence of 2.5 mM TCEP reducing agent. Thermal denaturation curves were plotted by the change of ellipticity at the global curve minimum and fitted with the sigmoidal two-state model to determine the melting temperature (T_m_) using Prism 8 (GraphPad).

### NMR

NMR experiments were performed as described previously (*9*). Briefly, proteins enriched in ^15^N were purified and buffer-exchanged to 10 mM sodium phosphate buffer containing 50 mM sodium chloride at pH 7, with 10% D_2_O. Proteins were concentrated to 300–500 μM. ^1^H,^15^N HSQC spectra were acquired in a 18.8T (800 MHz proton Larmor frequency) Bruker spectrometer equipped with a CPTC ^1^H,^13^C,^15^N 5 mm cryoprobe and an Avance III console, using 128 increments in the indirect dimension and a relaxation delay of 1 second.

### Negative-stain transmission electron microscopy

#### Sample preparation

4H.01 protein was incubated with Fabs (Motavizumab and 2D10) with equal molar ratios at 4 °C. After overnight incubation, complexes were purified on a Superose 6 Increase 10/300 column using an Äkta Pure system (GE Healthcare) in PBS buffer. Purified complexes were deposited at approximately 0.003 mg/ml onto carbon-coated copper grids (EMS, Hatfield, PA, United States) for 2 mins. Followed by a wash step with deionized water, the sample was stained with freshly prepared 0.75% uranyl formate for 4 mins.

#### Data acquisition

Digital images were collected using a direct detector camera Falcon III (Thermo Fisher) under an F20 electron microscope (Thermo Fisher) operated at 200 kV with the set-up of 4098 × 4098 pixels. The homogeneity and coverage of staining samples on the grid was first visualized at low magnification mode before automatic data collection. Automatic data collection was performed using EPU software (Thermo Fisher) at a nominal magnification of 50,000x, resulting in a pixel size of 2 Å, and defocus range from −1 μm to −2 μm.

#### Image processing

CTFFIND4 program (*54*) was used to estimate the contrast transfer function for each collected image. Around 1000 particles were manually selected using the installed package XMIPP within SCIPION framework (*55*). Manually picked particles served as input for XMIPP auto-picking utility, resulting in at least 10,000 particles. Selected particles were extracted with the box size of 100 pixels and subjected for three rounds of reference-free 2D classification without CTF correction using RELION-3.0 Beta suite (*56*).

### X-ray crystallization and structural determination

#### Co-crystallization of complex structure 101F Fab with 4E1H.95 and complex structure of Mota fab with 4H.01

The purified designed proteins were mixed with its target Fab in a 2:1 molar ratio (Mota in a complex of 4H.01 and 101F complexed with 4E1H.95) for overnight complex formation at 4 °C. The complex was purified by size exclusion chromatography (Hiload 16/600 Superdex 75 pg column, GE) in low ionic strength buffer (10 mM Tris, 50 mM NaCl, pH 8). The complex was subsequently concentrated to 10-12 mg/ml. Crystallization screening was performed in multiple 96-condition spare matrix suites available from Qiagen or Hampton Research. Crystals were grown at 291K using the sitting-drop vapor-diffusion method containing 100 nL of protein mixed with equal volume of precipitant. The 4E1H.95/101F Fab complex crystalized at 0.1 M sodium citrate pH 5.8, 0.1 M ammonium sulfate, 16% (w/v) PEG 4000, 20% (v/v) glycerol. The 4H.01/Mota Fab complex crystalized at 0.1 M bicine pH 9.3 with 22% (v/v) PEG Smear Broad. Crystals were flash-frozen in liquid nitrogen without adding cryo-protectant reagent.

#### Data collection and structural determination

Diffraction datasets were recorded at Paul Scherrer Institute with an X06DA (PXIII) beamline. The diffracted crystals of 4E1H.95/101F complex belongs to space group C121 and the 4H.01/Mota complex to the space group P3121. The diffraction data were integrated and processed to 2.7 Å (4E1H.95/101F) and 2.8 Å (4H.01/Mota) with X-ray Detector Software (XDS) (*57*). The structure was determined by the molecular replacement method using the PHASER module in PHENIX (*58, 59*). For the 4E1H.95/101F complex, the search of an initial phase was performed using a modelled complex of 4E1H.95 design model with the 101F Fab structure (PDB 3O41), which was obtained by superimposition of the design to the RSV fusion protein peptide. The initial search of 4H.01/Mota complex was done by using the Mota Fab structure (PDB 3IXT) and the computational model of 4H.01 protein, yielding clear molecular replacement solutions. Structural models were built and refined in iterative cycles using COOT (*60*) and PHENIX REFINE (*59*). The final refinement statistics are summarized in Tables S7 and S8.

### LUMABS sensor expression and characterization

#### Expression and purification

Genes encoding LUMABS were synthesized by Genscript, and cloned into a pET28 E.*coli* expression vector as described previously (*42*). For expression, constructs were transformed in E*.coli* BL21 (DE3) cells, grown until the OD reached 0.6, and induced with 1 mM IPTG. After overnight expression at 20 °C, cells were harvested, lysed and purified using NiNTA affinity chromatography and Strep-Tactin purification, followed by size-exclusion chromatography on a Superdex 200 increase column (GE Healthcare) using PBS as running buffer. Sensors were stored at −80 °C until further use.

#### Sensor characterization

LUMABS sensors were characterized as described previously (14). Briefly, the titrations with purified monoclonal antibody were performed in a total reaction volume of 20 μL, in white polystyrene 384 wells plates (Thermo Scientific™ Nunc™ 384-Well Polystyrene White Microplate, non-sterile, non-treated, 262360), on a Tecan Spark 10M. A sensor concentration of 50 pM or 100 pM was used in PBS supplemented with 1 mg/mL bovine serum albumin. After incubating for 45 minutes, NanoGlo assay substrate (Promega) was added (2000x dilution). Emission was subsequently recorded using a luminescence scan between 398 nm and 653 nm, with an integration time of 100 ms, at room temperature. For the characterization of the sensor in the presence of human plasma, pooled human plasma was spiked with purified monoclonal antibody and subsequently diluted by PBS supplemented with 1 mg/mL bovine serum albumin. Titrations were performed as described above, except for increased NanoGlo assay substrate concentration (1000x dilution) and integration time (1000 ms).

### Cell expression and testing of synthetic receptors

Human embryonic kidney cells (HEK-293T) were cultured in Dulbecco’s modified Eagle’s medium (DMEM, Life Technologies) supplemented with 10% (v/v) fetal bovine serum (FBS, Sigma-Aldrich) and 1% (v/v) streptomycin/penicillin at 37 °C and 5% CO_2_. For cell-based experiments, 1.5 × 10^6^ cells diluted in 12 mL DMEM were seeded into 96-well plates 24 h before transfection. The transfection mixture for 6 wells consisted of 800 ng plasmid DNA (300 ng each of the receptor plasmids pLeo1336 and pLeo1340 or 600 ng for single receptor controls, 100 ng of the TetR-Elk1 fusion protein plasmid mKP37, and 10 ng of the TetR reporter plasmid pTS1017) was added into 300 μL DMEM and mixed with 4 μg of polyethyleneimine (Polysciences Inc.). The transfection mixture was vortexed and incubated for 20 min before adding into the inner 60 wells of the cell culture plate. Medium was exchanged 15h post-transfection for 125 μL/well medium with serial dilutions of inducer and inhibitor as indicated in the respective figures (Fig. 5). Supernatants were analysed for quantification of the secreted reporter protein SEAP after 24 h. The data was analyzed with Prism 8 (GraphPad) and the response curve was fitted to a dose-response model to determine the EC_50_ and the shift of EC_50_ in the presence of inhibitor.

