## Supplementary material for "A bottom-up approach for the *de novo* design of functional proteins": Supp material

### Supplementary information

#### Supplementary Figures

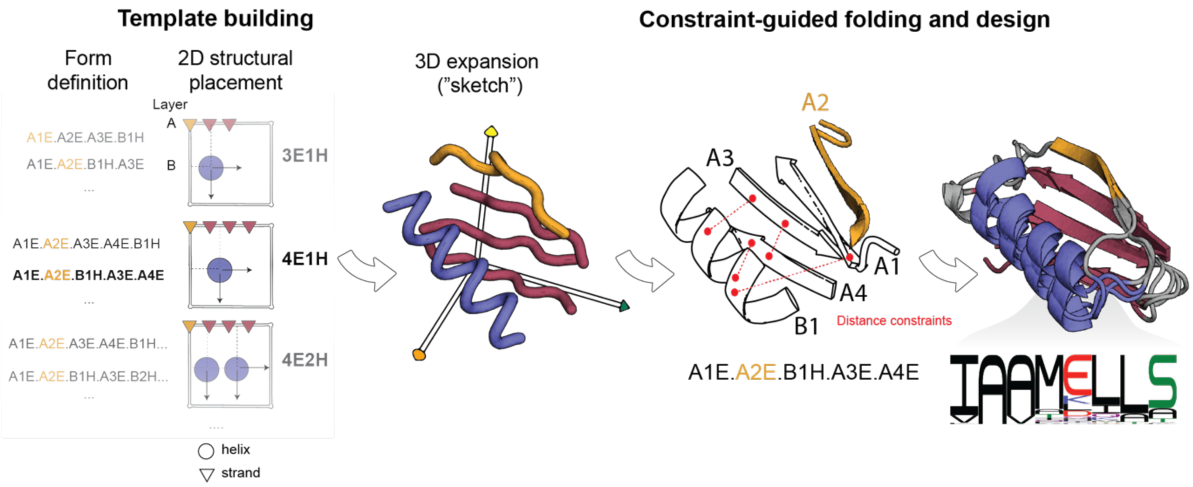


**Fig. S1**
**Integrated pipeline for the generation of *de novo* backbones with the TopoBuilder followed by conformational sampling and sequence design.** The TopoBuilder design pipeline consists of two steps, the building of idealized design templates followed by a folding and design step using Rosetta FunFolDes. Protein topologies are first described in a string-based format, allowing to quickly enumerate possible protein topologies that can be assembled around a given motif (orange). Selected forms are then expanded in the 3D space, yielding an idealized sketch. Atom-pair and distance constraints are derived from the sketch, which subsequently guide a folding and sequence design protocol (Rosetta FunFolDes).

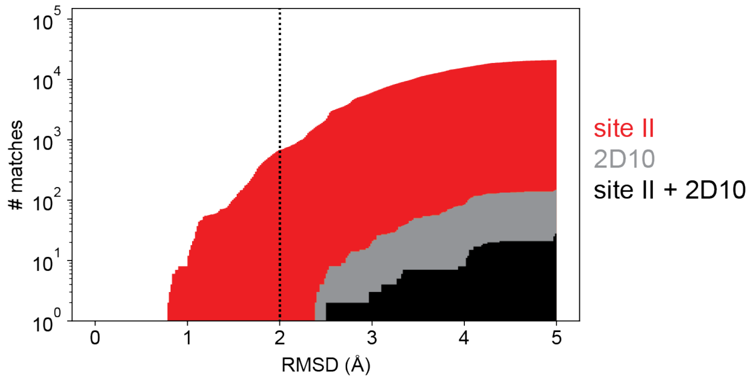


**Fig. S2**
**Design templates that accommodate both the RSVF site II and the RSVG 2D10 epitope are sparse in the natural protein repertoire.** We used Rosetta MotifGraft (*1*) to search the PDB (03/2014) for structural templates with local structural similarity to site II (PDB 3IXT, red) and 2D10 (PDB 5WN9, grey). Proteins that can accommodate both motifs in non-overlapping regions are shown in black. The number of unique matches is shown on the y-axis, as a function of the C-alpha RMSD to the query motif. Templates with an RMSD of < 2 Å to a query motif are often amenable for grafting methods, as indicated by a dotted black line.

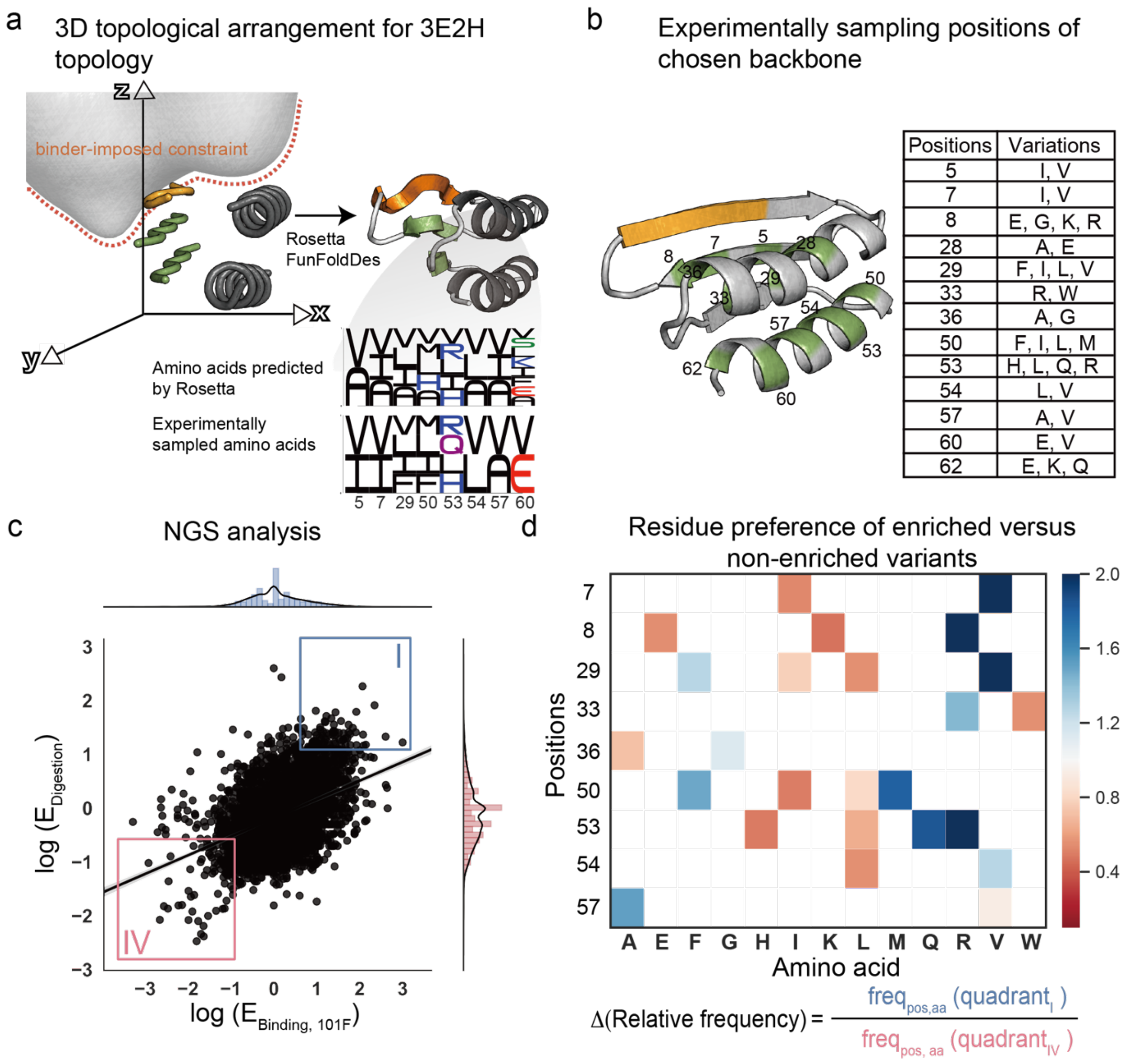


**Fig. S3**

**Computational design, experimental screening and enrichment analysis of the 3E2H design series. a**, TopoBuilder assembly of the 3E2H topology (left) and a full atom model after folding and design (right). The logo plots show the sequence diversity in selected core positions as predicted by Rosetta FastDesign and the diversity for each position encoded in a combinatorial library (using degenerate codons) for yeast display screening. **b**, Detailed view on all positions experimentally sampled (green). **c**, Enrichment analysis following next-generation sequencing (NGS) of populations sorted for high affinity binding (x-axis) versus resistance to protease digestion (y-axis). **d**, Residue preferences for each position when comparing sequences positively enriched for both binding and protease resistance (c, quadrant I, blue) versus sequences that were negatively enriched (c, quadrant IV, red). 100-200 sequences each were analyzed. The heatmap shows the relative frequency of the respective amino acids in quadrant I versus quadrant IV, showing, for example, an overrepresentation of valine over isoleucine in position 7 in sequences from quadrant I.

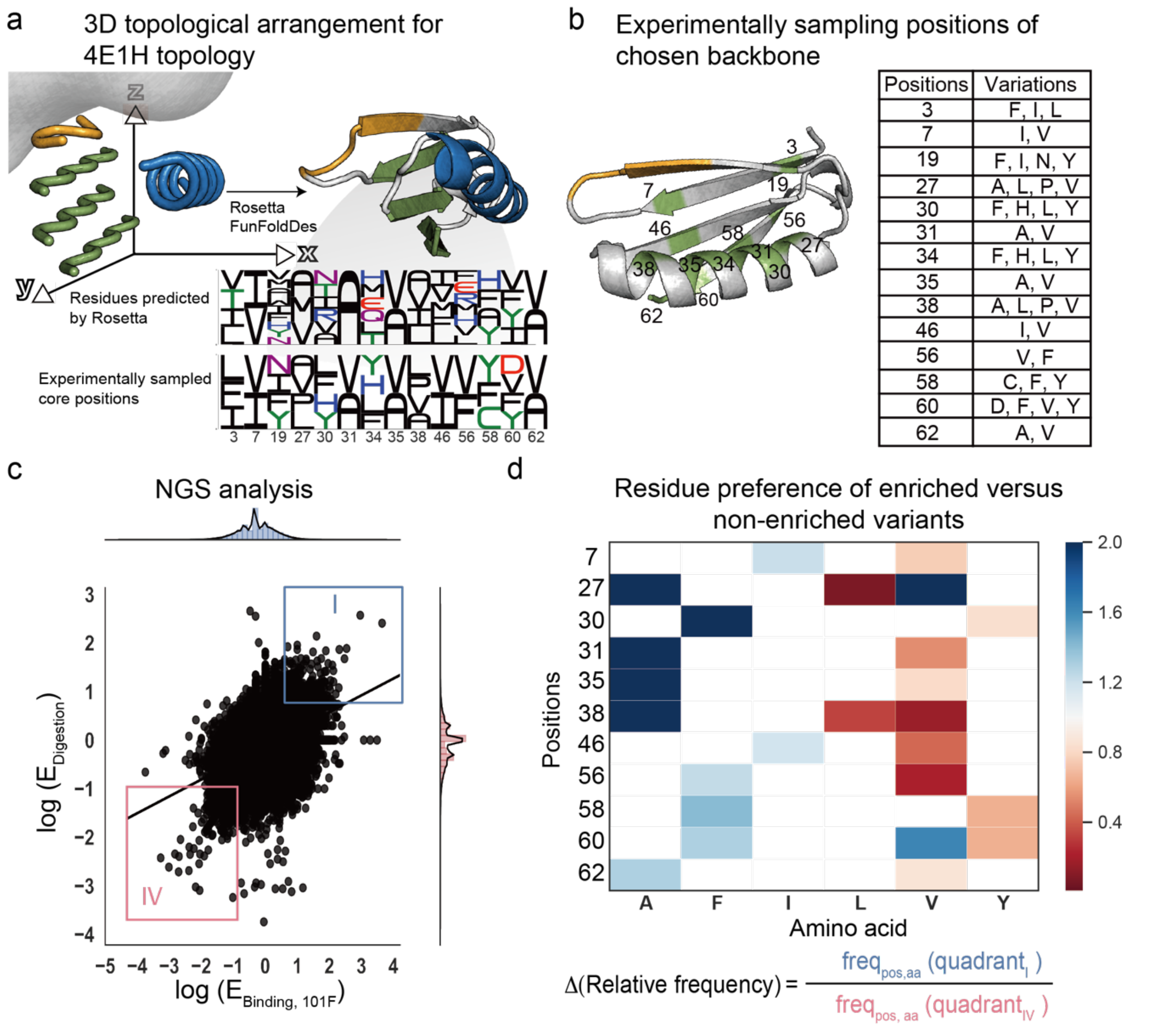


| Positions | Variations |
|---|---|
| 3 | F, I, L |
| 7 | I, V |
| 19 | F, I, N, Y |
| 27 | A, L, P, V |
| 30 | F, H, L, Y |
| 31 | A, V |
| 34 | F, H, L, Y |
| 35 | A, V |
| 38 | A, L, P, V |
| 46 | I, V |
| 56 | V, F |
| 58 | C, F, Y |
| 60 | D, F, V, Y |
| 62 | A, V |

**Fig. S4**

**Computational design, experimental screening and enrichment analysis of the 4E1H design series**. **a**, TopoBuilder assembly of the 4E1H topology. **b**, Detailed view on all positions experimentally sampled (green). **c**, Enrichment analysis of dual selection pressures, binding to 101F antibody and resistance to protease digestion. **d**, Residue preferences for indicated position in positively enriched versus negatively enriched sequences. See Fig. S3 caption for details.

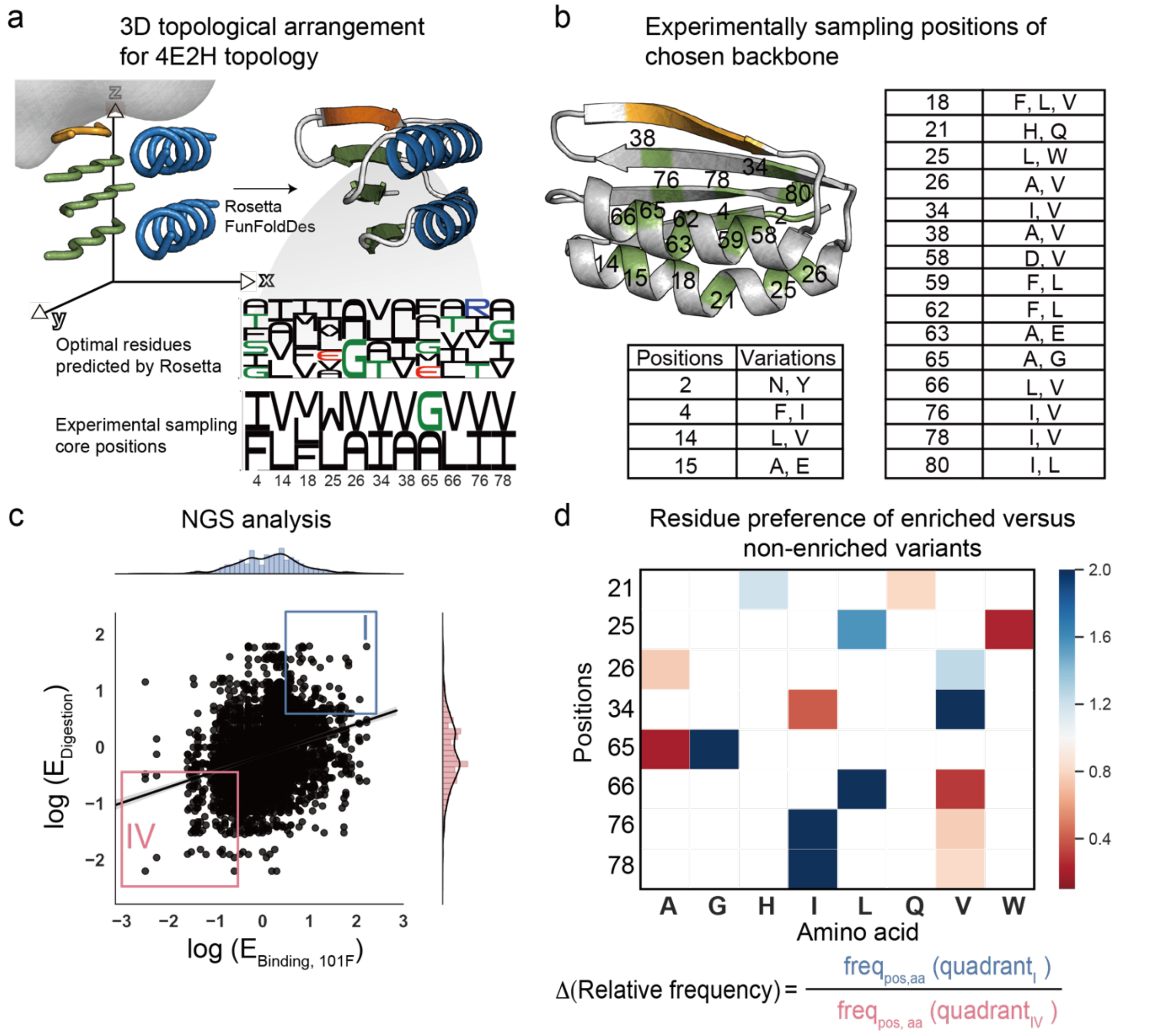


**Fig. S5**

**Computational design, experimental screening and enrichment analysis of the 4E2H design series. a**, TopoBuilder assembly of the 4E2H topology and folded full-atom structure after Rosetta FunFoldDes. **b**, Detailed view on all positions experimentally sampled (green). **c**, Enrichment analysis. **d**, Residue preferences for indicated position in positively enriched versus negatively enriched sequences. See Fig. S3 caption for details.

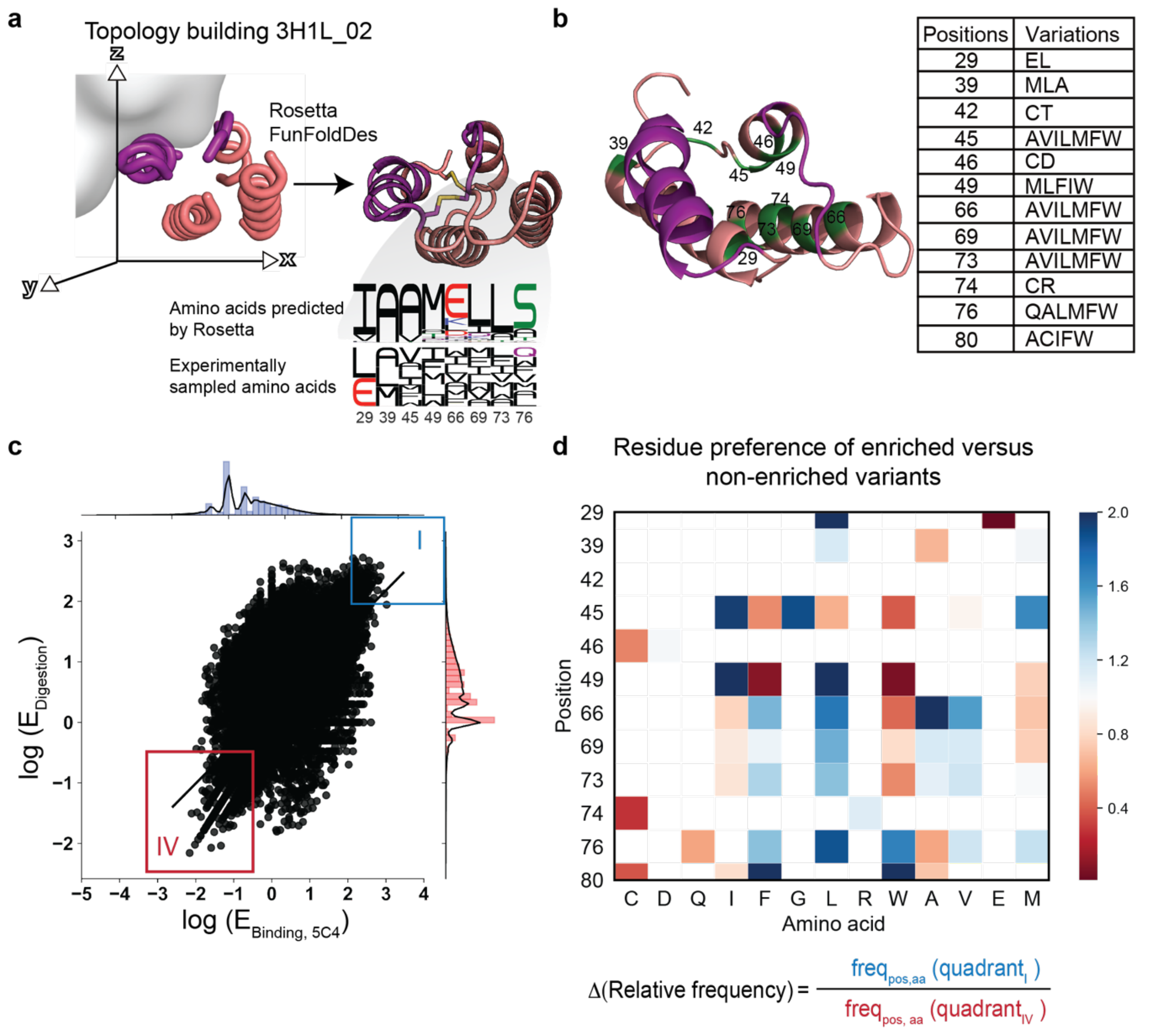


**Fig. S6**

**Computational design, experimental screening and enrichment analysis of the 3H1L_02 design series.** **a**, TopoBuilder assembly of the 3H1L_02 topology and folded full-atom structure after Rosetta FunFoldDes, the epitope region is highlighted in purple. **b**, Detailed view on all positions experimentally sampled (green). **c**, Enrichment analysis. **d**, Residue preferences for indicated position in positively enriched versus negatively enriched sequences. See Fig. S3 caption for details.

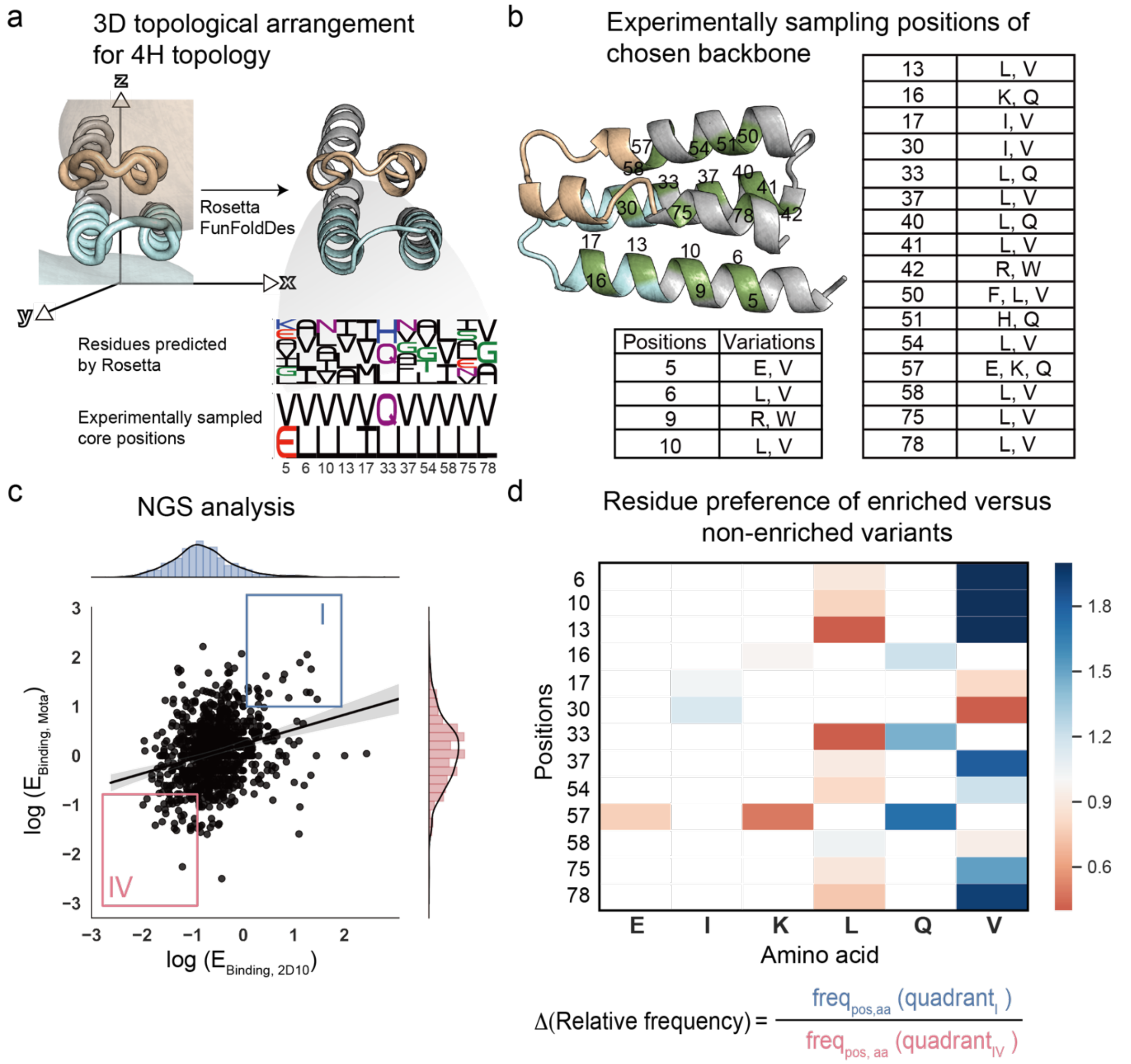


**Fig. S7**

**Computational design, experimental screening and enrichment analysis of the 4H design series. a**, TopoBuilder assembly of the 4H topology and folded full-atom structure after Rosetta FunFoldDes, the Mota epitope region is colored in cyan and 2D10 epitope is colored in wheat. **b**, Detailed view on all positions experimentally sampled (green). **c**, Enrichment analysis. **d**, Residue preferences for indicated position in positively enriched versus negatively enriched sequences. See Fig. S3 caption for details.

**a**

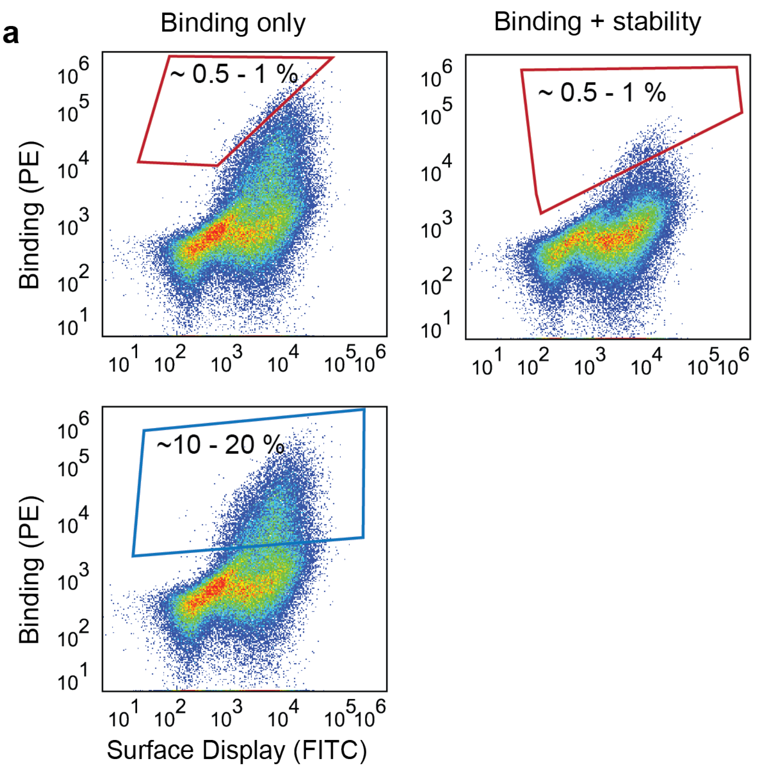


**b**

$$\text{Enrichment (E)} = \frac{\text{freq}_{\text{Seq}}\ \text{(high selection pressure)}}{\text{freq}_{\text{Seq}}\ \text{(low selection pressure)}}$$

**Fig. S8**

**Gating strategy and enrichment analysis. a**, Yeast libraries were sorted under two selective pressures (binding, binding following protease treatment to digest partially unfolded designs), and the best 0.5 - 1% of clones were sorted (red sorting gates). To normalize the frequency of each sequence, we sorted the library under a low selection pressure (binding only, 10 - 20%, blue sorting gate). **b**, Enrichment scores were computed for each sequence for binding and stability. The score was calculated as the relative frequency of each sequence under high selective pressure versus low selective pressure.

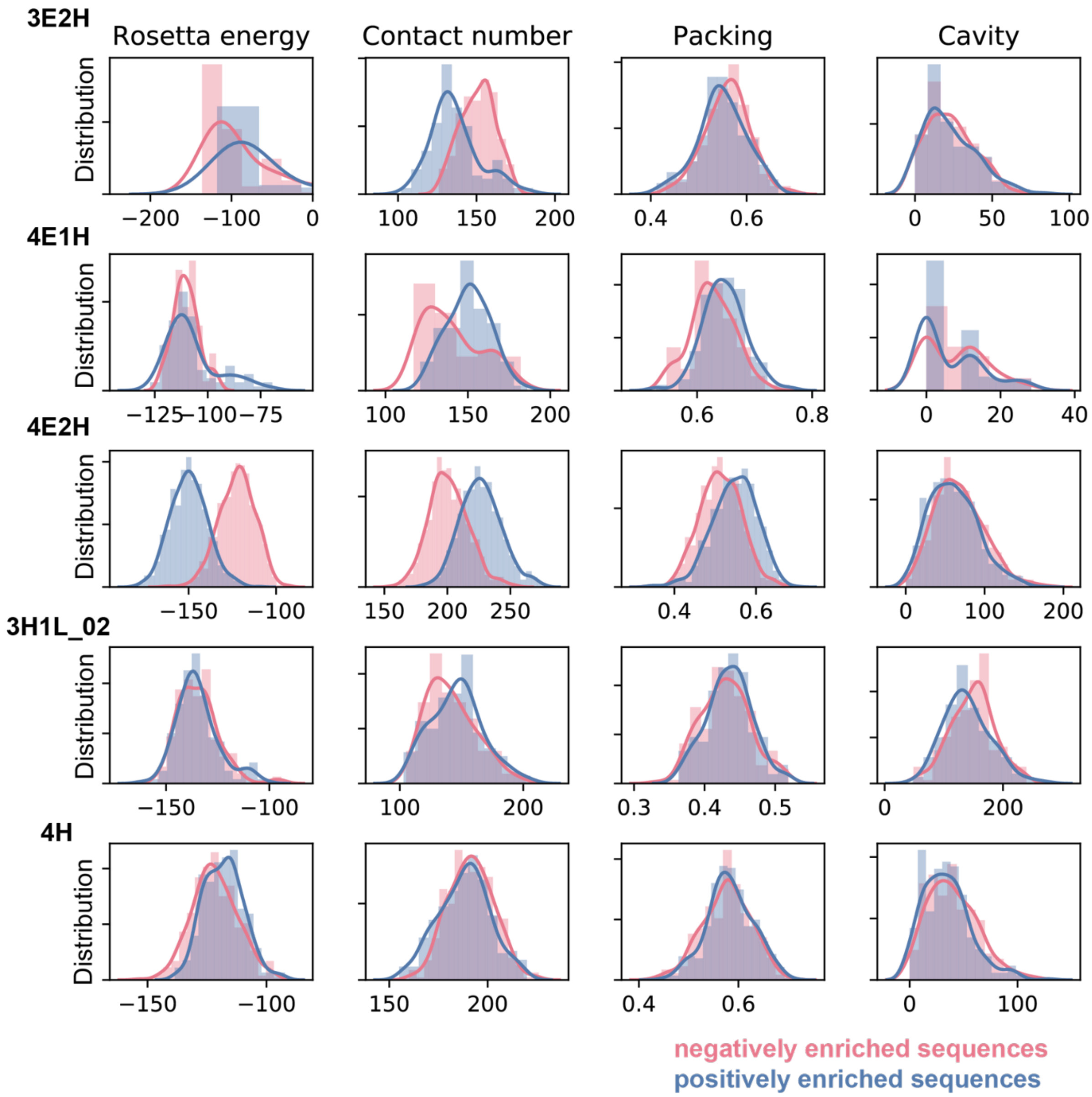


**Fig. S9**

**Comparison of structural metrics for computational models of positively and negatively enriched sequences from different folds.** For each topology, computational models were generated using Rosetta FastDesign for 100-200 sequences with the strongest negative (red) and the strongest positive enrichment (blue), respectively. The distribution of four different scores (total Rosetta energy, number of side-chain carbon-carbon contacts in the protein core with a distance cutoff of 4.5 Å (contact number), packing score and the volume of intra-protein voids in Å$^3$ (cavity)) is shown for negatively and positively enriched populations.

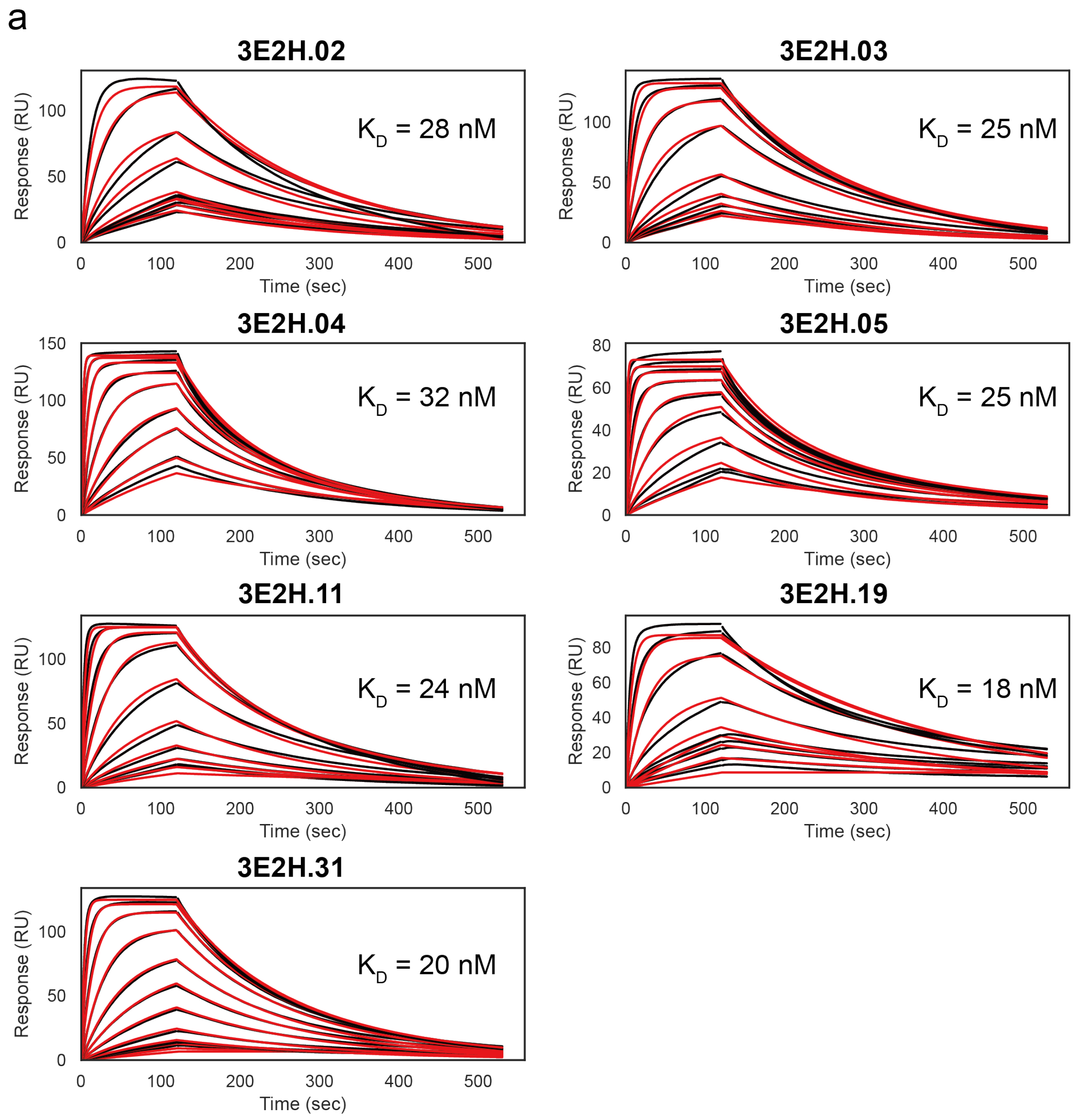


*Continued next page*

b

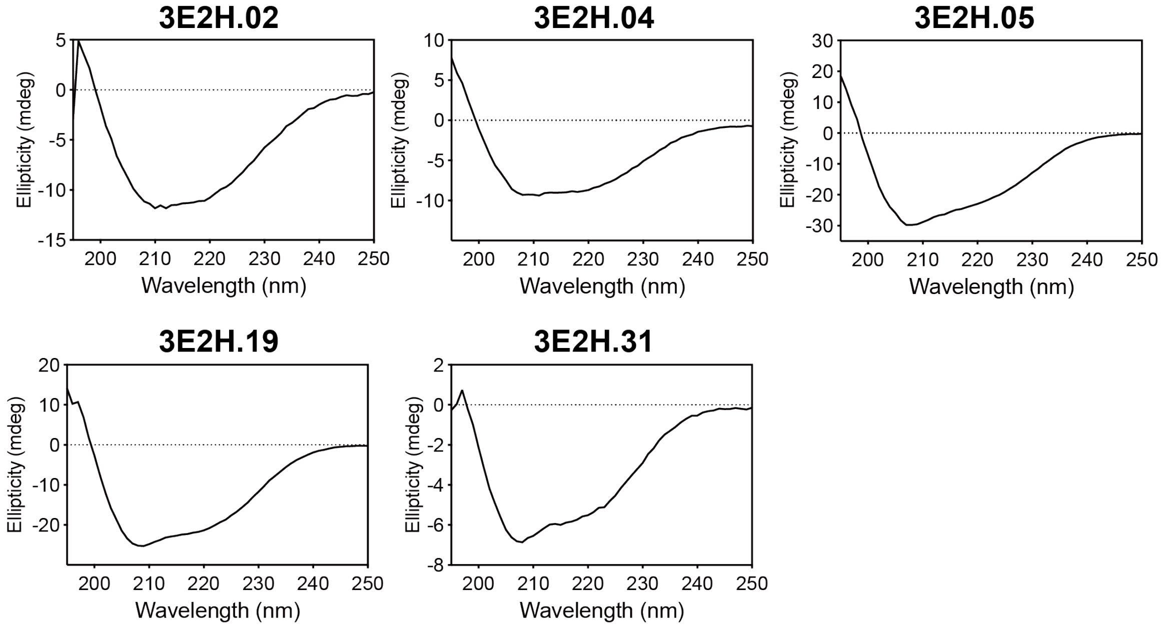


c

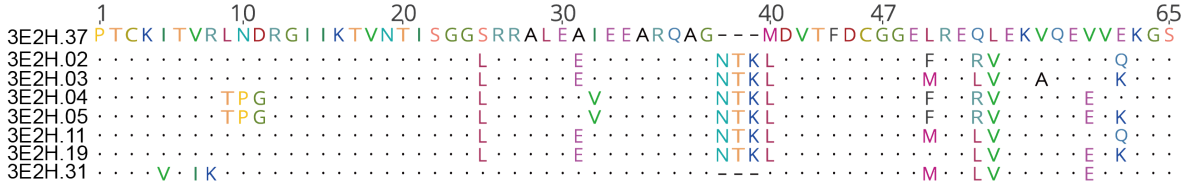


**Fig. S10**

**Biophysical characterization of 3E2H design series. a**, The binding affinities of selected 3E2H designs was measured by SPR. The 3E2H designs were immobilized on the sensor chip surface, flowing 101F Fab as analyte. Kinetic dissociation constants ($K_D$s) were obtained by fitting the curves using a 1:1 Langmuir model. Raw data are shown in black, fitted curves in red. **b**, CD spectra of selected designs at 20 °C. **c**, Sequence comparison between biophysically characterized clones. The pairwise sequence identity between the selected 8 sequences is 88%.

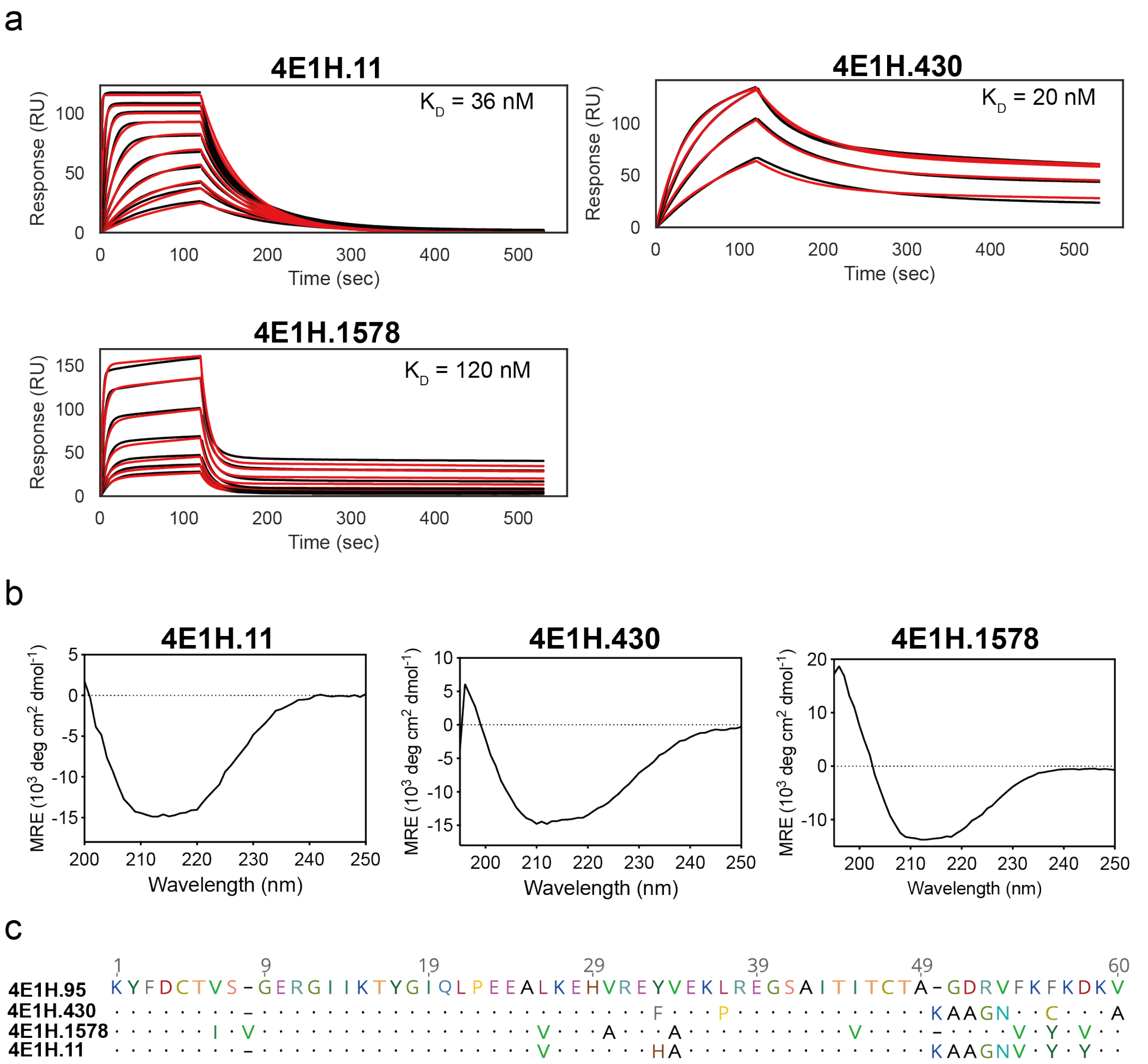


**Fig. S11**

**Biophysical characterization of 4E1H design series. a**, Binding affinities of selected 4E1H designs to 101F. 101F IgG was immobilized on the sensor chip surface, flowing 4E1H designs as analyte. Curves were fitted using a 1:1 Langmuir model, yielding $K_D$s ranging between 20 nM and 120 nM. Raw data are shown in black, fitted curves in red. **b**, CD spectra of selected designs at 20 °C. **c**, Sequence comparison between enriched clones. The pairwise sequence identity between the 4 selected sequences is 82%.

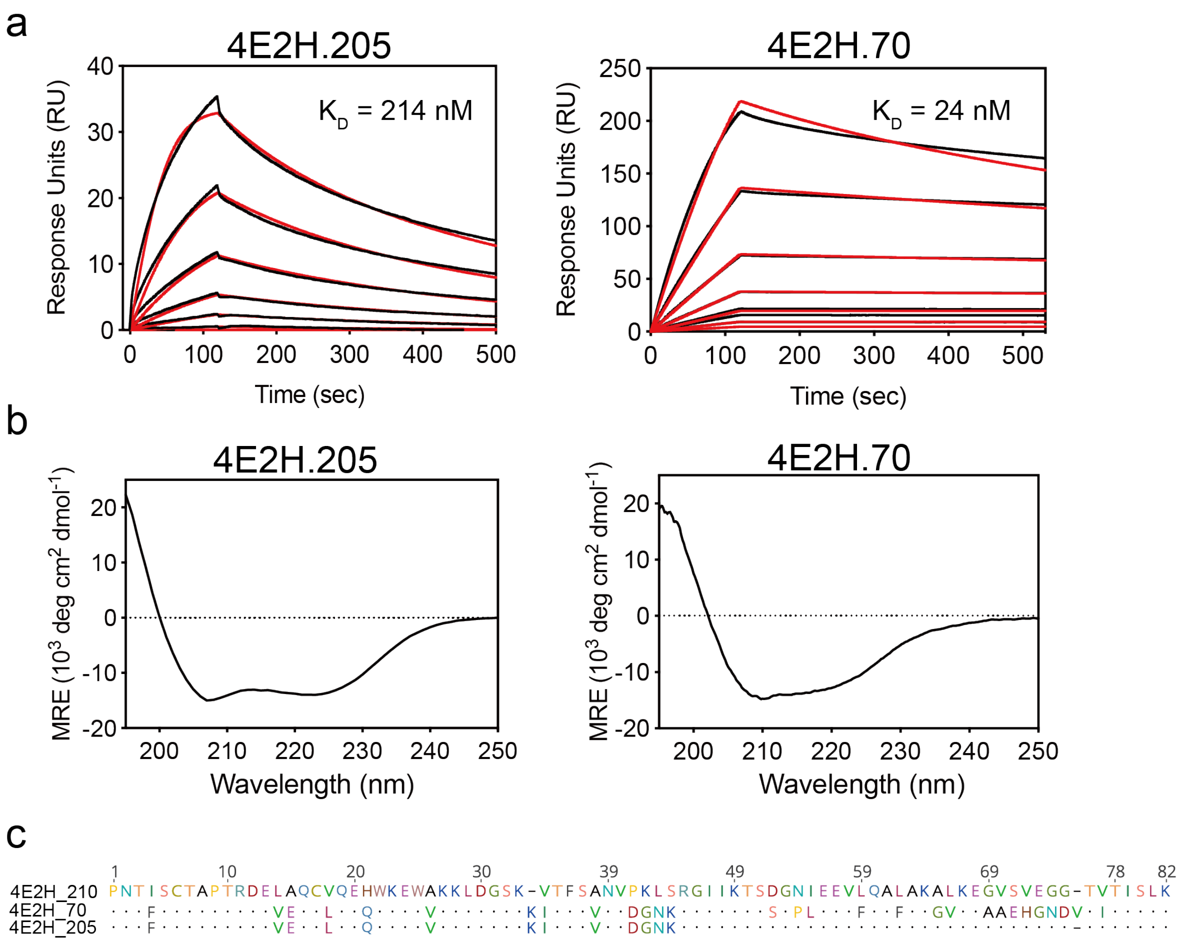


**Fig. S12**

**Biophysical characterization of 4E2H design series. a**, The binding affinities of selected 4E2H designs to 101F. 101F IgG was immobilized as ligand on the sensor chip surface, flowing the designs as analyte. Raw data are shown in black, fitted curves in red. **b**, CD spectra of selected 4E2H designs at 20 °C. **c**, Sequence comparison between enriched clones. The pairwise sequence identity between the selected designs is around 75%

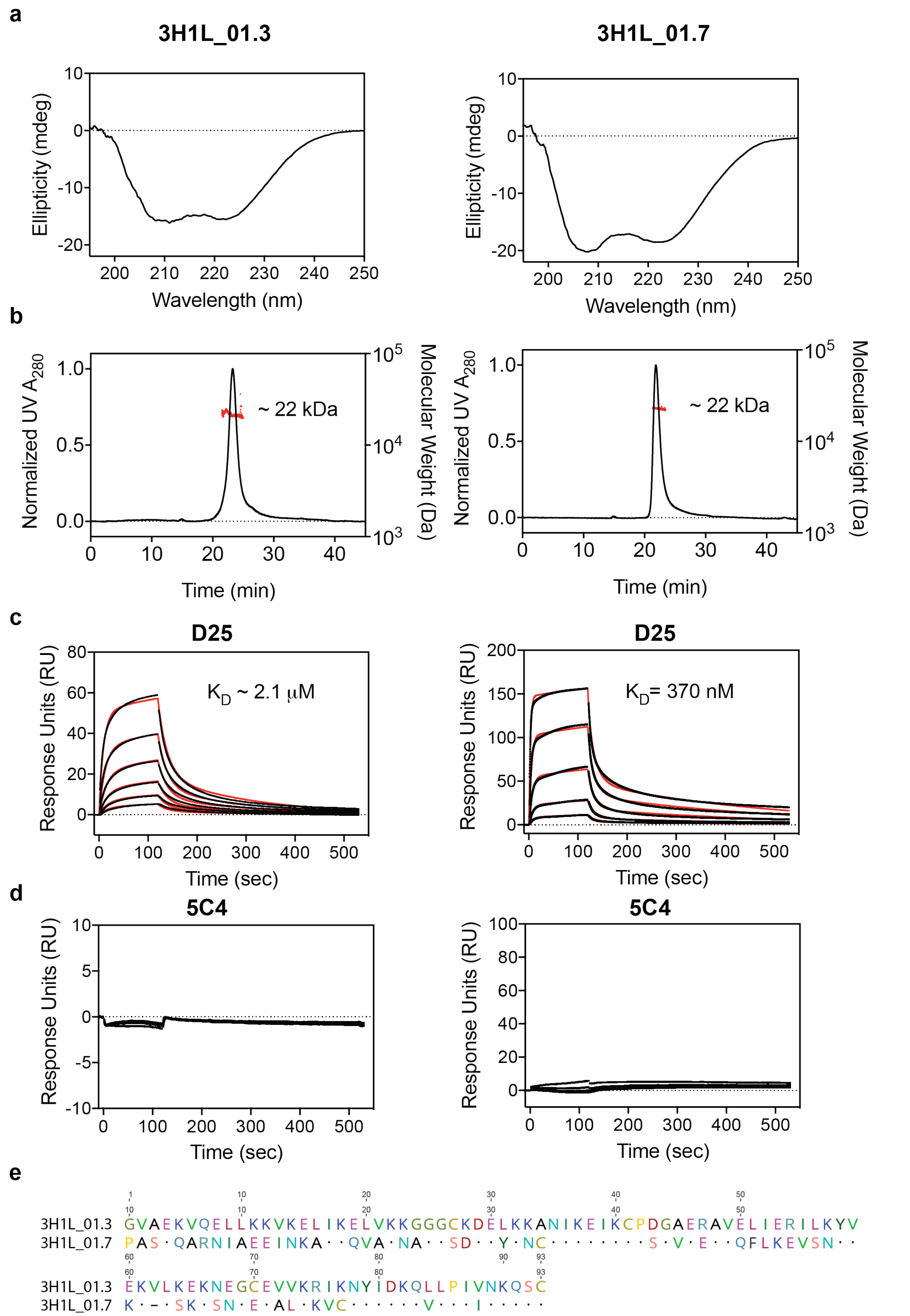


**Fig. S13**

**Biophysical characterization of 3H1L_01 designs. a**, CD spectra at 25 °C indicate that proteins adopt alpha helical conformations in solution. **b**, SEC-MALS profiles of 3H1L_01 designs show the formation of dimers in solution (molecular weight ~22 kDa, theoretical monomer molecular weight 11.7 and 11.5 kDa, for 3H1L_01.3 and 3H1L_01.7, respectively). **c-d**, Measurement of binding affinity to D25 (c) and 5C4 (d) by SPR. 3H1L_01 designs were immobilized on the sensor chip surface, and D25 or 5C4 Fabs were injected as analyte in various concentrations. Kinetic dissociation constants are indicated, with no detectable binding to 5C4. Raw data are shown in black, fitted curves in red. **e**, The pairwise sequence identity between the two designs shown is 45%.

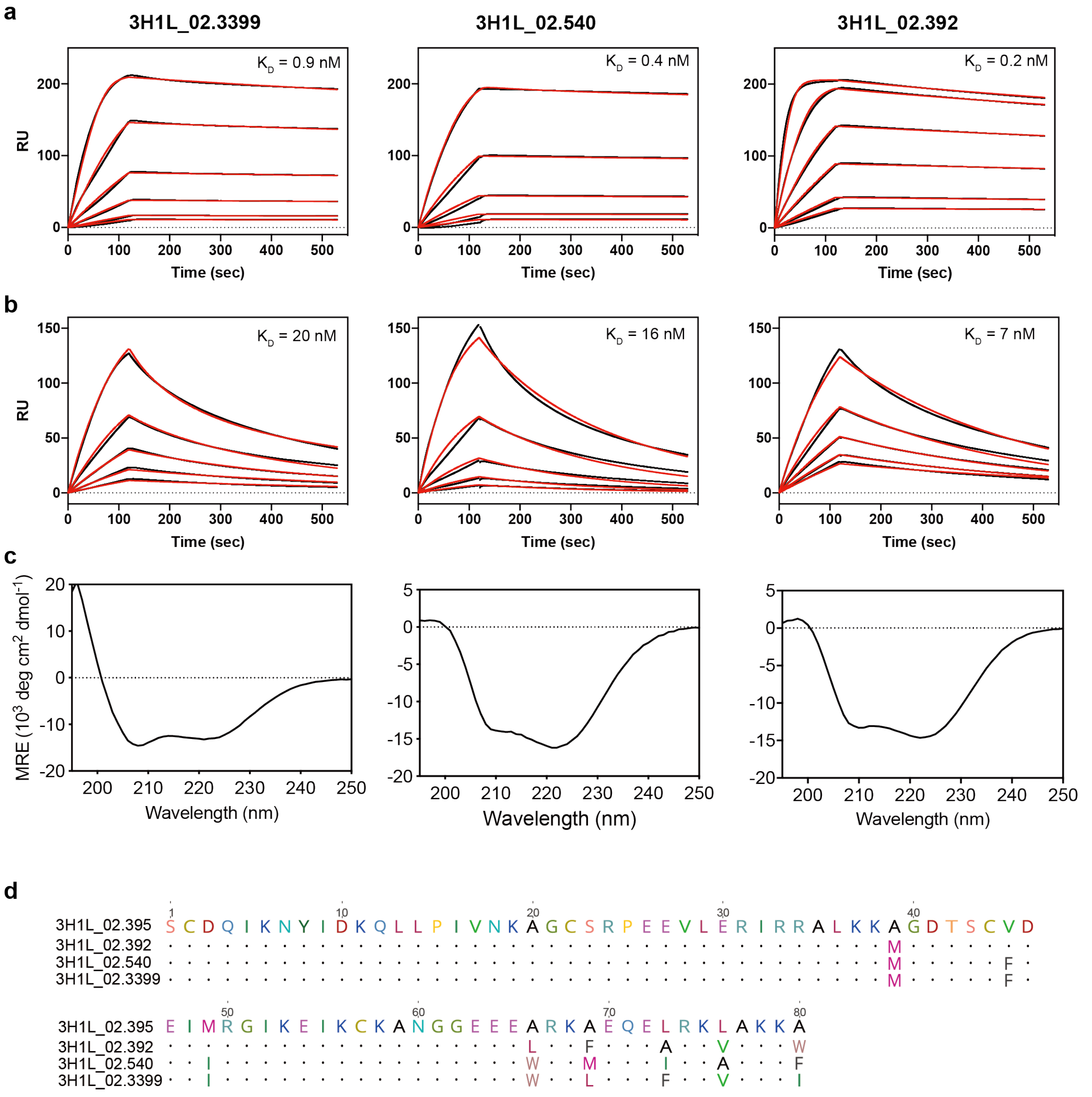


**Fig. S14**

**Biophysical characterization of 3H1L_02 designs. a**, SPR binding curves of 3H1L_02.3399, 3H1L_02.540 and 3H1L_02.392 to D25. D25 IgG was immobilized on the sensor chip surface, flowing 3H1L_02 variants as analyte. **b**, Binding of 3H1L_02 variants to 5C4 IgG. Raw data are shown in black, fitted curves in red. All data were fitted with a 1:1 Langmuir binding model. **c**, CD spectra at 25 °C indicate a predominantly helical secondary structure content. **d**, Sequence comparison between 4 enriched clones. The pairwise sequence identity between the selected sequences is 92%.

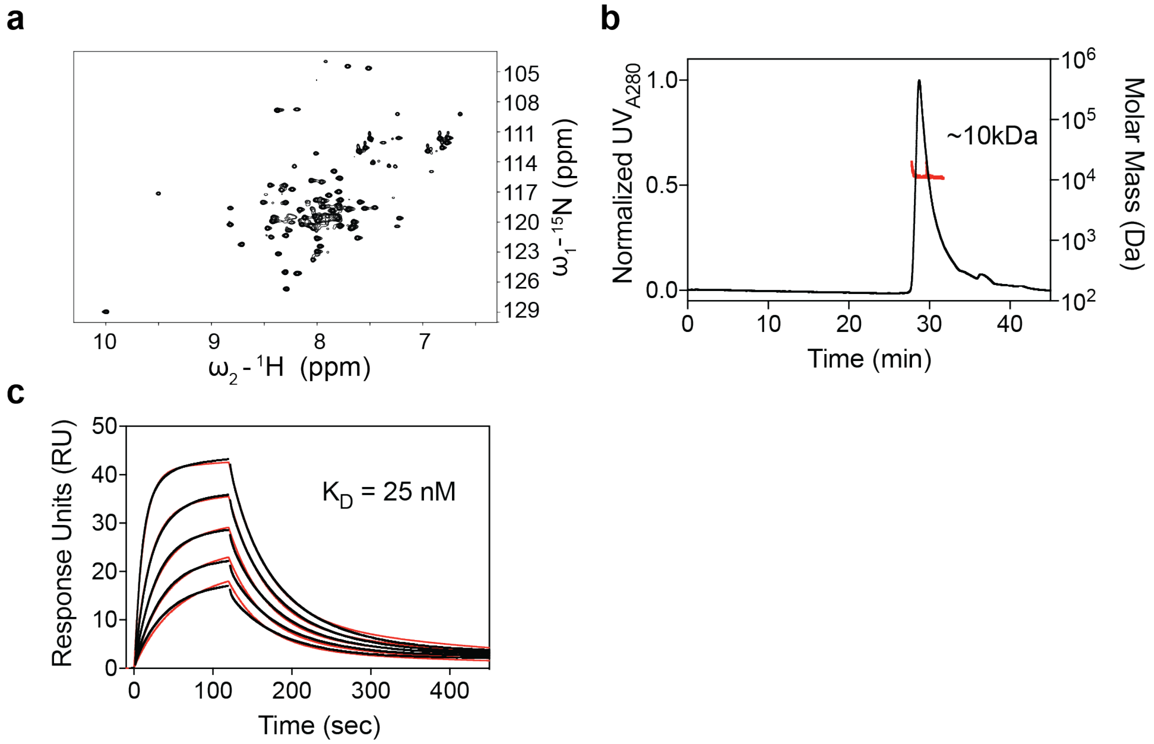


**Fig. S15**

**Extended biophysical characterization of 3H1L_02.395. a**, The 2D NMR $^{15}N$ HSQC spectrum for 3H1L_02.395 is well dispersed, confirming that it is well-folded in solution. **b**, 3H1L_02.395 is monomeric with a determined molecular weight of 10 kDa closely matching its theoretical molecular weight of 10.5 kDa. **c**, 3H1L_02.395 binds with a $K_D$ of 25 nM to 5C4, as determined by SPR. Raw data are shown in black, curve fits (1:1 Langmuir model) are shown in red.

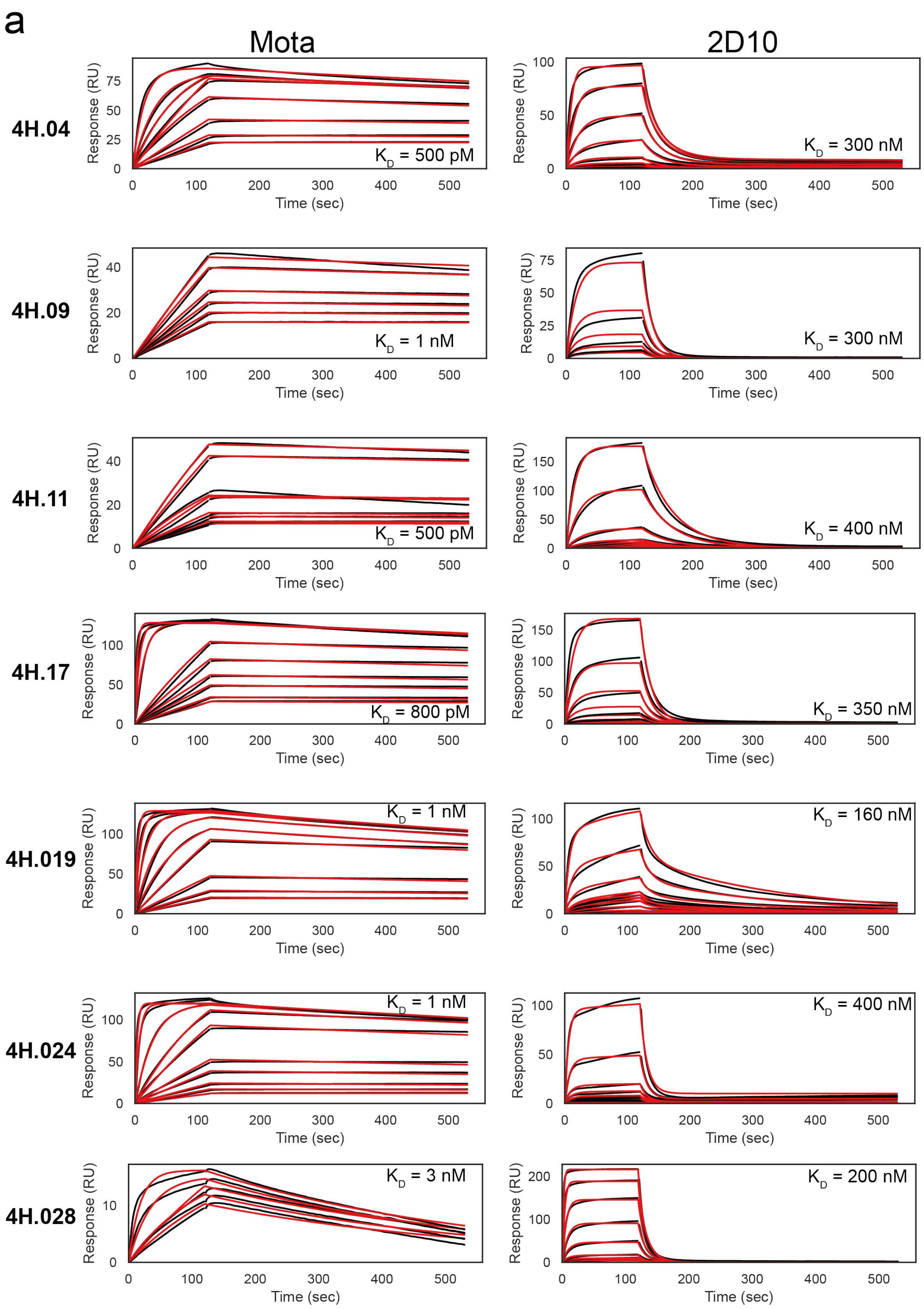


*Continued next page*

b

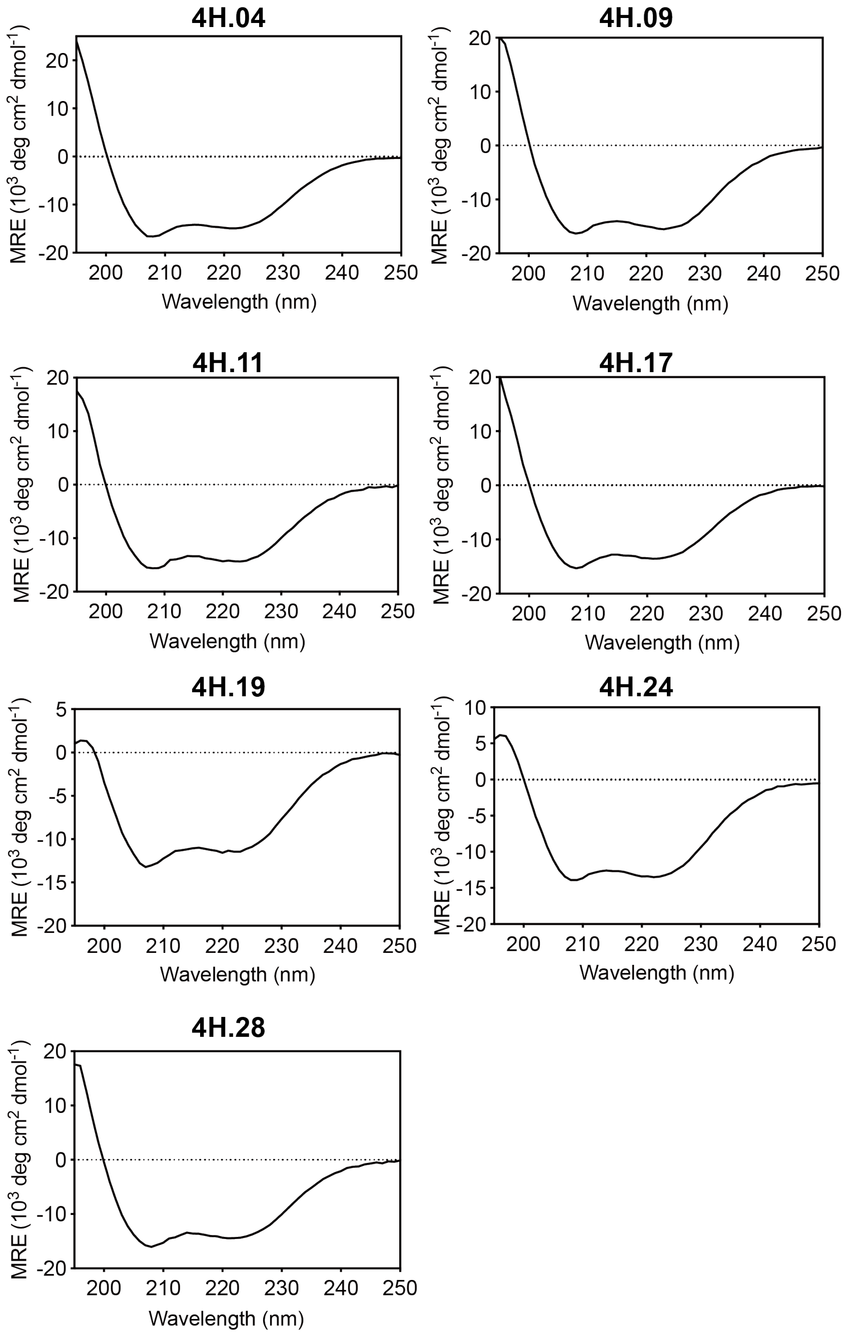


c

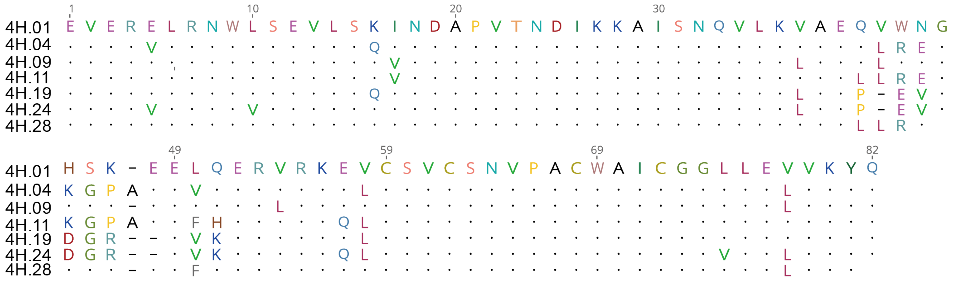


**Fig. S16**

**Biophysical characterization of 4H designs. a**, SPR binding curves of 4H designs to motavizumab (Mota, left) and 2D10 (right). IgGs were immobilized on the sensor chip surface, flowing 4H variants as analyte. Raw data and curve fits (1:1 Langmuir kinetic fit) are shown in black and red, respectively. **b**, CD spectra at 25 °C indicate a predominantly helical secondary structure content. **c**, Sequence comparison between enriched clones. The pairwise sequence identity between the selected 7 sequences is 85%.

a

**4H.01**

b

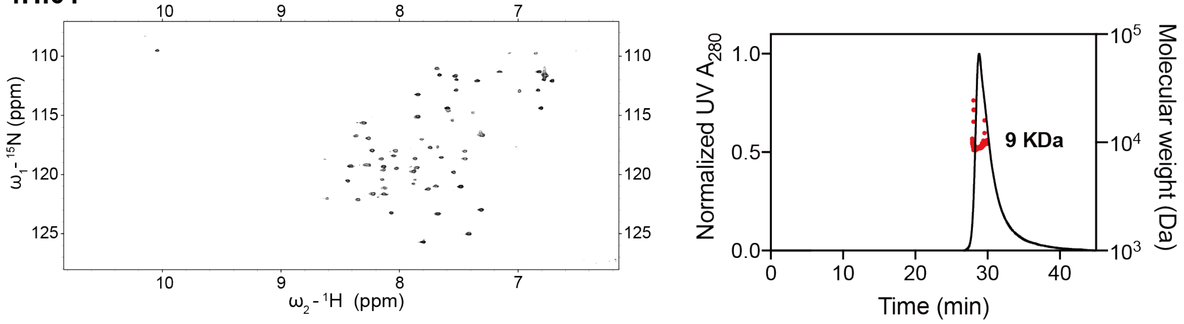


**Fig. S17**

**Extended biophysical characterization of 4H.01. a**, 2D NMR $^{15}N$ HSQC spectrum for 4H.01 is well dispersed, confirming that it is well-folded in solution. **b**, SEC-MALS profile for 4H.01 confirms that it is monomeric in solution with a measured molecular weight in solution of 9 kDa, which is in close agreement with the theoretical monomer mass of 9.9 kDa.

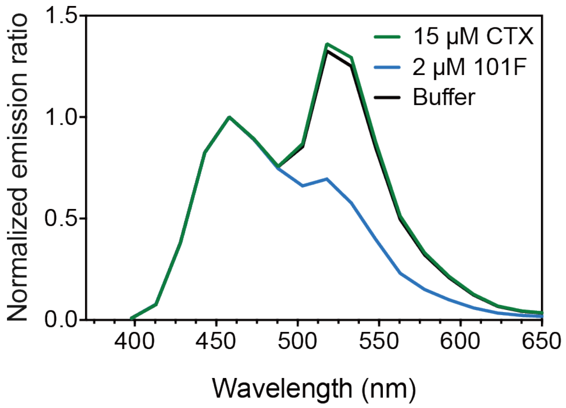


**Fig. S18**

**Specificity of LUMABS sensor for the target antibody.** Luminescence spectra of the 4E2H.210 LUMABS sensor in the absence of antibody (black), compared to 2 μM 101F IgG (blue) or 15 μM cetuximab (CTX, green), an anti-EGFR antibody, showing that the sensor only responds in the presence of epitope-specific antibodies.

### Supplementary Tables

| | | sequence distance (#mutations) | | | |
|---|---|---|---|---|---|
| Design | core position sampled | minimum | maximum | average | average core identity (%) |
| 3E2H | 8 | 2 | 5 | 3.5 | 56 |
| 4E1H | 14 | 5 | 10 | 7.5 | 46 |
| 4E2H | 11 | 4 | 10 | 6.2 | 44 |
| 3H1L_02 | 9 | 2 | 6 | 4.6 | 49 |
| 4H | 11 | 6 | 8 | 6.5 | 41 |

**Table S1**
Sequence distance between computationally designed sequences and sequences enriched under two selective pressures. The topologies built by the TopoBuilder were folded and designed using Rosetta FunFolDes, and filtered for the best 1,000 sequences with lowest Rosetta energy. Sequences that were enriched under two selective pressures in the yeast screening (150 - 200 sequences with strongest enrichments) were compared against the computationally designed sequences (sampled core positions only). Between 8 and 14 core positions were sampled experimentally, and the closest Rosetta designs differed from 2 to 6 mutations, with an average ranging from 3.5 to 7.5 mutations. The average identity in the core residues was 47%, ranging between 41% and 56% sequence identity.

| Design | E value (blast) | Dali match (closest RMSD, Å) |
|---|---|---|
| 3E2H.37 | 0.25 | 2.5 |
| 4E1H.95 | 3.6 | 1.9 |
| 4E2H.210 | 2.1 | 2.4 |
| 3H1L_02.395 | 0.09 | 2.4 |
| 4H.01 | 1.00E-06 | 2.6 |

**Table S2**
Sequence and structure comparison of designed sequences to the natural repertoire. The protein sequences of selected designs were blasted against a database of non-redundant protein sequence in NCBI, with the resulting E values indicating that the designed sequences are dissimilar to naturally occurring proteins. To compare the designs structurally, a Dali search (*2*) was performed against the PDB, and the closest RMSD is reported.

**Table S3**: Primers used for constructing combinatorial sequence libraries for 4E1H, 3E2H, 4E2H and 4H.

| Primer name | Sequence |
|---|---|
| **4E1H** | |
| 4E1H_O1 | GACAATAGCTCGACGATTGAAGGTAGATACCCATACGACGTTCCAGACTACGCTCTGCAGGCTAGTGGTGGAGGAGG |
| 4E1H_O2 | CCCTCCGCCTCCGCTACCGCCTCCACCAGAGCCTCCTCCACCACTAGCCTG |
| 4E1H_O3.1 | CGGTAGCGGAGGCGGAGGGTCGGCTAGCCATATGAAGTATHTCGACTGTACGRTCAGCGGGGAGCGCGGCATCATCAAGACCTAC |
| 4E1H_O3.2 | CGGTAGCGGAGGCGGAGGGTCGGCTAGCCATATGAAGTATHTCGACTGTACGRTCAGCGTGGGGGAGCGCGGCATCATCAAGACCTAC |
| 4E1H_O4.1 | CGGGGGATCCCTCACGARSTTTTTCARCGWRCTCACGARCGWRTTCCTTARSGGCCTCATGGCTCCCCTCGTCGWWTCCGTAGGTCTTGATGATGCCGCG |

| | |
|---|---|
| 4E1H_O4.2 | CGGGGGATCCCTCACGARSTTTTTCARCGWRCTCACGARCGWRTTCCTTARSGGCCTCCTCAGGAAGCTGGWWTCCGTAGGTCTTGATGATGCCGCG |
| 4E1H_O4.3 | GGGATCCCTCACGARSTTTTTCARCGWRCTCACGARCGWRTTCCTTARSGGCCTCATGATCAATATCGGGGTGGWWTCCGTAGGTCTTGATGATGCCGCG |
| 4E1H_O5.1 | GGCGGATCCGCGATTACCRTCACCTGCACTGCCAAAGCTGCCGGTAATKTCAAATDCAAGKWCAAGGYTGGCTCGTGGGGCTCGCACC |
| 4E1H_O5.2 | GGCGGATCCGCGATTACCRTCACCTGCACTGCCGGTGACCGTGTGKTCAAATDCAAGKWCAAGGYTGGCTCGTGGGGCTCGCACC |
| 4E1H_O5.3 | GGCGGATCCGCGATTACCRTCACCTGCACTGGCGGTGGCTTACGTKTCAAATDCAAGKWCAAGGYTGGCTCGTGGGGCTCGCACC |
| 4E1H_O6 | CAGAAATAAGCTTTTGTTCGGACCCGGGCTCAGCCTATTAGTGGTGGTGGTGGTGGTGCGAGCCCCACGAGCC |
| 4E1H_O7 | GGGTCCGAACAAAAGCTTATTTCTGAAGAGGACTTGTAATAGCTCGAGATCTGATAAC |
| 4E1H_O8 | GTACGAGCTAAAAGTACAGTGGGAACAAAGTCGATTTTGTTACATCTACACTGTTGTTATCAGATCTCGAGCTATTACAAGTCC |
| **3E2H** | |
| 3E2H_O1 | GACAATAGCTCGACGATTGAAGGTAGATACCCATACGACGTTCCAGACTACGCTCTGCAGGCTAGTGGTGGAGGAGGC |
| 3E2H_O2 | CATATGGCTAGCCGACCCTCCGCCTCCGCTACCGCCTCCACCAGAGCCTCCTCCACCACTAGCCTGCAG |
| 3E2H_O3.1 | CGGAGGGTCGGCTAGCCATATGCCGACCTGCAAGRTTACGRTCRRGTTGGAGGAACGCGGGATCATCAAGACGGTC |
| 3E2H_O3.2 | CGGAGGGTCGGCTAGCCATATGCCGACCTGCAAGRTTACGRTCRRGACACCCGGTCGCGGGATCATCAAGACGGTC |
| 3E2H_O3.3 | CGGAGGGTCGGCTAGCCATATGCCGACCTGCAAGRTTACGRTCRRGCTGAATGATCGCGGGATCATCAAGACGGTC |
| 3E2H_O4.1 | GCCGCAATCAAAGGTCACGTCCAACTTCGTATTASCGGCCTGCCRTGCCTCCTCAANCKCCTCTAAGGCACGATGATCGGTAGTGTTGACCGTCTTGATGATCCCGCG |
| 3E2H_O4.2 | GCCGCAATCAAAGGTCACGTCGCCCGGCGCACCASCGGCCTGCCRTGCCTCCTCAANCKCCTCTAAGGCACGCTTTGACCCGGTTCCGTTGACCGTCTTGATGATCCCGCG |
| 3E2H_O4.3 | GCCGCAATCAAAGGTCACGTCCAACTTCGTATTASCGGCCTGCCRTGCCTCCTCAANCKCCTCTAAGGCACGGCGCGAGCCGCCACTGATGGTGTTGACCGTCTTGATGATCCCGCG |
| 3E2H_O5 | GACGTGACCTTTGATTGCGGCGGTGAAWTSCGCGAGCDVSTGGAGAAAGYGCAAGAAGWAGTCVAGAAGGGCTCGTGGGGCTCGCAC |
| 3E2H_O6 | AGAAATAAGCTTTTGTTCGGATCCGGGCTCAGCCTATTAGTGGTGGTGGTGGTGGTGCGAGCCCCACGAGCC |
| 3E2H_O7 | GAGCCCGGATCCGAACAAAAGCTTATTTCTGAAGAGGACTTGTAATAGCTCGAGATCTGATAACAACAGTGTAGATGTAACAAAATCGAC |
| 3E2H_O8 | GTACGAGCTAAAAGTACAGTGGGAACAAAGTCGATTTTGTTACATCTACACTGTTGTTATCAGATCTC |
| **4E2H** | |
| 4E2H_O1 | GACAATAGCTCGACGATTGAAGGTAGATACCCATACGACGTTCCAGACTACGCTCTGCAGGCTAGTG |
| 4E2H_O2 | CCGCTACCGCCTCCACCAGAGCCTCCTCCACCACTAGCCTGCAGAGCGTAGTC |
| 4E2H_O3 | GGTGGAGGCGGTAGCGGAGGCGGAGGGTCGGCTAGCCATATG |
| 4E2H_O4.1 | TGGCGCTGCCGACGGATCCGACTTTCTTARCCMATTCTTTCCAWTGTTCCTGAAVGCATTGCKCAASTTCGTCTGGCACGTCCATAGTGCAGCTGAWAGTGTWTGGCATATGGCTAGCCGACCCTCC |
| 4E2H_O4.2 | TGGCGCTGCCGACGGATCCGTCTAATTTCTTARCCMATTCTTTCCAWTGTTCCTGAAVGCATTGCKCAASTTCGTCGCGAGTGGGAGCAGTGCAGCTGAWAGTGTWTGGCATATGGCTAGCCGACCCTCC |

| | |
|---|---|
| 4E2H_O4.3 | TGGCGCTGCCGACGGATCCGCCTTCCTTARCCMATTCTTTCCAWTGTTCCTGAAVGCATTGCKCAASTTCGTCCTTGGAGACAACAGTGCAGCTGAWAGTGTWTGGCATATGGCTAGCCGACCCTCC |
| 4E2H_O5.1 | GGCTATCGTCACACCGGATCCAAAAAARTCACTTTTAGCGYTAACGTAGACGGGAACAAACGCGGCATCATCAAGACC |
| 4E2H_O5.2 | GGCTATCGTCACACCGGATCCAAAGGARTCACTTTTAGCGYTAACGTGCCTAAACTGTCACGCGGCATCATCAAGACC |
| 4E2H_O5.3 | GGCTATCGTCACACCGGATCCAAARTCACTTTTAGCGYTAACGTGCCTAAACTGTCACGCGGCATCATCAAGACC |
| 4E2H_O5.4 | GGCTATCGTCACACCGGATCCAAAGGARTCACTTTTAGCGYTAACGACCCGAAAGAGATCCGCGGCATCATCAAGACC |
| 4E2H_O6.1 | GGTGCGAGCCCCACGAGCCCTTTAWGCTGAYAGTGAYTGTAACGAAACCAGAAGCGCCTTCCTTAASASCTTTCKCAARTGCCTGAARAWCCTCCTCGATGTTTCCGTCGGAGGTCTTGATGATGCCGCG |
| 4E2H_O6.2 | GGTGCGAGCCCCACGAGCCCTTTAWGCTGAYAGTGAYTGTGCCTCCTTCAACTGATACCCCTTCCTTAASASCTTTCKCAARTGCCTGAARAWCCTCCTCAAGAGGCCCCGACGAGGTCTTGATGATGCCGCG |
| 4E2H_O6.3 | GGTGCGAGCCCCACGAGCCCTTTAWGCTGAYAGTGAYTGTTACATCATTACCATGTTCGGCGGCTTCCTTAASASCTTTCKCAARTGCCTGAARAWCCTCCTCAAGAGGCCCCGACGAGGTCTTGATGATGCCGCG |
| 4E2H_O7 | GGCTCGTGGGGCTCGCACCACCACCACCACCACTAATAGGCTGAGCCCGGGTCC |
| 4E2H_O8 | CTCGAGCTATTACAAGTCCTCTTCAGAAATAAGCTTTTGTTCGGACCCGGGCTCAGCCTATTAG |
| 4E2H_O9 | CTGAAGAGGACTTGTAATAGCTCGAGATCTGATAACAACAGTGTAGATGTAACAAAATCGACTTTG |
| 4E2H_O10 | GTACGAGCTAAAAGTACAGTGGGAACAAAGTCGATTTTGTTACATCTACACTGTTG |
| **4H** | |
| 4H_O1 | GACAATAGCTCGACGATTGAAGGTAGATACCCATACGACGTTCCAGACTACGCTCTGCAGGCTAGTGGTGGAGGAGGC |
| 4H_O2 | CATATGGCTAGCCGACCCTCCGCCTCCGCTACCGCCTCCACCAGAGCCTCCTCCACCACTAGCCTGCAG |
| 4H_O3 | CGGAGGGTCGGCTAGCCATATGGAAGTTGAACGCGWASTTCGCAACYGGSTTTCGGAASTTTTGAGTMAGRTCAACGATGCACCAGTGACCAACGACATC |
| 4H_O4.1 | CACGTTGGAGCAGACGCTACAAASCTBTTTACGCASACGCTCWTGWAMCTCCTCTTTGCTGTGACCATTCCRAASCWGTTCTGCAASTTTTAACACCWGGTTCGAAAYGGCCTTCTTGATGTCGTTGGTCACTGGTGCATCG |
| 4H_O4.2 | CACGTTGGAGCAGACGCTACAAASCTBTTTACGCASACGCTCWTGWAMCTCCTCGGCTGGCCCTTTCCCCTCCCRAASCWGTTCTGCAASTTTTAACACCWGGTTCGAAAYGGCCTTCTTGATGTCGTTGGTCACTGGTGCATCG |
| 4H_O4.3 | CACGTTGGAGCAGACGCTACAAASCTBTTTACGCASACGCTCWTGWAMCTCCTCGTCCGCACCGCCCGTCCRAASCWGTTCTGCAASTTTTAACACCWGGTTCGAAAYGGCCTTCTTGATGTCGTTGGTCACTGGTGCATCG |
| 4H_O4.4 | CACGTTGGAGCAGACGCTACAAASCTBTTTACGCASACGCTCCTTAACTTCGCGTCCGTCTCCGACTTCTGGTTCTGCAASTTTTAACACCWGGTTCGAAAYGGCCTTCTTGATGTCGTTGGTCACTGGTGCATCG |
| 4H_O5 | TGTAGCGTCTGCTCCAACGTGCCCGCGTGCTGGGCGATTTGTGGAGGTSTTTTAGAASTTGTAAAGTATCAAGGCTCGTGGGGCTCGCACCAC |
| 4H_O6 | GGATCCGGGCTCAGCCTATTAGTGGTGGTGGTGGTGGTGCGAGCCCCACGAGCC |
| 4H_O7 | CACTAATAGGCTGAGCCCGGATCCGAACAAAAGCTTATTTCTGAAGAGGACTTGTAATAGCTCGAGATCTGATAACAA |
| 4H_O8 | GTACGAGCTAAAAGTACAGTGGGAACAAAGTCGATTTTGTTACATCTACACTGTTGTTATCAGATCTCGAGCTATTACAAGTCCTC |

**Table S4**: Combinatorial library for 3H1L_02 topology as assembled using a defined set of codons.

| **Template sequence** | |
|---|---|
| TCCTGTGACCAGATAAAGAATTACATAGATAAGCAGCTTTTGCCTATTGTAAACAAAGCCGGATGCAGTAGACCGGAGGAAGTAGAAGAGAGGATTAGACGTGCATTAAAAAAAATGGGCGATACAAGTTGCTTCGACGAAATAATGAGAGGCATCAAGGAGATTAAGTGTAAGGCGAATGGTGGAGAGGAGGAGATGCGTAAGATGGAGCAAGAGGCCAGGAAACAGGCTAAGAAGGCG | |
| **Position** | **Alternative codons** |
| 29 | TTA |
| 39 | TTA,GCC |
| 42 | TGC |
| 45 | GGC, GTT, ATC, TTA, ATG, TGG |
| 46 | TGC |
| 49 | TTA, TTC, ATC, TGG |
| 66 | GCC, GTT, TTA, TTC, ATC, TGG |
| 69 | GCC, GTT, TTA, TTC, ATC, TGG |
| 73 | ATG, GTT, TTA, TTC, ATC, TGG |
| 74 | TGC |
| 76 | GCC, GTT, TTA, ATG, TTC, TGG |
| 80 | TGC, ATC, TTC, TGG |

**Table S5**: Amino acid sequences of designed proteins tested

| **Designed topology** | **Sequence** |
|---|---|
| **4E1H** | |
| 4E1H.11 | KYFDCTVSGERGIIKTYGIQLPEEAVKEHVREHAEKLREGSAITITCTAKAAGNVKYKYKVGSWGS |
| 4E1H.95 | KYFDCTVSGERGIIKTYGIQLPEEALKEHVREYVEKLREGSAITITCTAGDRVFKFKDKVGSWGS |
| 4E1H.430 | KYFDCTVSGERGIIKTYGIQLPEEALKEHVREFVEKPREGSAITITCTAKAAGNFKCKDKAWGS |
| 4E1H.1578 | KYFDCTISVGERGIIKTYGIQLPEEAVKEHAREYAEKLREGSAITVTCTAGDRVVKYKVKVWGS |
| **3E2H** | |
| 3E2H.02 | PTCKITVRLNDRGIIKTVNTISGGLRRALEEIEEARQAGNTKLDVTFDCGGEFRERVEKVQEVVQKGS |
| 3E2H.03 | PTCKITVRLNDRGIIKTVNTISGGLRRALEEIEEARQAGNTKLDVTFDCGGEMRELVEKAQEVVKKGS |
| 3E2H.04 | PTCKITVRTPGRGIIKTVNTISGGLRRALEAVEEARQAGNTKLDVTFDCGGEFRERVEKVQEEVEKGS |
| 3E2H.05 | PTCKITVRTPGRGIIKTVNTISGGLRRALEAVEEARQAGNTKLDVTFDCGGEFRERVEKVQEEVKKG |

| | |
|---|---|
| 3E2H.11 | PTCKITVRLNDRGIIKTVNTISGGLRRALEEIEEARQAGNTKLDVTFDCGGEMRELVEKVQEVVQKGS |
| 3E2H.19 | PTCKITVRLNDRGIIKTVNTISGGLRRALEEIEEARQAGNTKLDVTFDCGGELRELVEKVQEEVKKGS |
| 3E2H.31 | PTCKVTIKLNDRGIIKTVNTISGGSRRALEAIEEARQAGMDVTFDCGGEMRELVEKVQEEVEKGS |
| 3E2H.37 | PTCKITVRLNDRGIIKTVNTISGGSRRALEAIEEARQAGMDVTFDCGGELREQLEKVQEVVEKGS |
| **4E2H** | |
| 4E2H.70 | PNTFSCTAPTRDEVEQCLQEQWKEWVKKLDGSKKITFSVNVDGNKRGIIKTSSGPLEEVFQAFAKGVKEAAEHGNDVTITISLKGS |
| 4E2H.205 | PNTFSCTAPTRDEVEQCLQEQWKEWVKKLDGSKKITFSVNVDGNKRGIIKTSDGNIEEVLQALAKALKEGVSVEGGTVTISLKGS |
| 4E2H.210 | PNTISCTAPTRDELAQCVQEHWKEWAKKLDGSKVTFSANVPKLSRGIIKTSDGNIEEVLQALAKALKEGVSVEGGTVTISLKGS |
| **3H1L_01** | |
| 3H1L_01.3 | GVAEKVQELLKKVKELIKELVKKGGGCKDELKKANIKEIKCPDGAERAVELIERILKYVEKVLKEKNEGCEVVKRIKNYIDKQLLPIVNKQSC |
| 3H1L_01.7 | GPASEQARNIAEEINKAIKQVAKNAGGSDDEYKNCNIKEIKCPSGVEEAVQFLKEVSNYVKKLSKKSNGEEALKKVCNYIDKQVLPIINKQSCL |
| **3H1L_02** | |
| 3H1L_02.129 | SCDQIKNYIDKQLLPIVNKAGCSRPEEVLERIRRALKKLGDTSCFDEIIRGIKEIKCKANGGEEEIRKVEQEFRKMAKKI |
| 3H1L_02.392 | SCDQIKNYIDKQLLPIVNKAGCSRPEEVLERIRRALKKMGDTSCVDEIMRGIKEIKCKANGGEEELRKFEQEARKVAKKW |
| 3H1L_02.395 | SCDQIKNYIDKQLLPIVNKAGCSRPEEVLERIRRALKKAGDTSCVDEIMRGIKEIKCKANGGEEEARKAEQELRKLAKKA |
| 3H1L_02.540 | SCDQIKNYIDKQLLPIVNKAGCSRPEEVLERIRRALKKMGDTSCFDEIIRGIKEIKCKANGGEEEWRKMEQEIRKAAKKF |
| 3H1L_02.901 | SCDQIKNYIDKQLLPIVNKAGCSRPEEVLERIRRALKKMGDTSCFDEILRGIKEIKCKANGGEEEMRKVEQEARKFAKKW |
| 3H1L_02.3399 | SCDQIKNYIDKQLLPIVNKAGCSRPEEVLERIRRALKKMGDTSCFDEIIRGIKEIKCKANGGEEEWRKLEQEFRKVAKKI |
| **4H** | |
| 4H.01 | EVERELRNWLSEVLSKINDAPVTNDIKKAISNQVLKVAEQVWNGHSKEELQERVRKEVCSVCSNVPACWAICGGLLEVVKYQ |
| 4H.04 | EVERVLRNWLSEVLSQINDAPVTNDIKKAISNQVLKVAEQLREGKGPAEEVQERVRKELCSVCSNVPACWAICGGLLELVKYQ |
| 4H.09 | EVERELRNWLSEVLSKVNDAPVTNDIKKAISNQVLKLAEQLWNGHSKEELQERLRKEVCSVCSNVPACWAICGGLLELVKYQ |
| 4H.11 | EVERELRNWLSEVLSKVNDAPVTNDIKKAISNQVLKVAELLREGKGPAEEFHERVRKQLCSVCSNVPACWAICGGLLEVVKYQ |
| 4H.17 | EVERVLRNWLSEVLSQINDAPVTNDIKKAISNQVLKLAEQLREGKGPAEEVQERVRKELCSVCSNVPACWAICGGLLELVKYQ |
| 4H.19 | EVERELRNWLSEVLSQINDAPVTNDIKKAISNQVLKLAEPEVGDGREVKERVRKELCSVCSNVPACWAICGGLLEVVKYQ |
| 4H.24 | EVERVLRNWVSEVLSKINDAPVTNDIKKAISNQVLKLAEPEVGDGREVKERVRKQLCSVCSNVPACWAICGGVLELVKYQ |
| 4H.28 | EVERELRNWLSEVLSKINDAPVTNDIKKAISNQVLKVAELLRNGHSKEEFQERVRKEVCSVCSNVPACWAICGGLLELVKYQ |

**Table S6**: Plasmids used for engineered cells and biosensors

| **Plasmid** | **Description** | **Experiment** | **Reference / genbank ID** |
|---|---|---|---|

| | | | |
|---|---|---|---|
| MKP37 | $P_{hCMV}$-TetR-ELK1-pA<br><br>Mammalian expression vector for the MAPK reporter protein TetR-ELK1 | Fig. 6<br><br>used in combination with a TetR reporter plasmid to monitor activity of the MAPK pathway | Keeley et. al (*3*) |
| pLeo1336 | $P_{SV40}$- ScFv_2D10-$EpoR_{ex}$-$FGFR1_{int}$-pA<br><br>Mammalian expression vector for a synthetic receptor binding to the 2D10 epitope of 4H.01 | Fig. 6<br><br>Activation of the MAPK pathway in response to 4H.01 | This work |
| pLeo1340 | $P_{SV40}$- ScFv_Mota-$EpoR_{ex}$-$FGFR1_{int}$-pA<br><br>Mammalian expression vector for a synthetic receptor binding to the Mota epitope of 4H.01 | Fig. 6<br><br>Activation of the MAPK pathway in response to 4H.01 | This work |
| pTS1017 | $P_{TetO7}$-SEAP-pA<br><br>Mammalian reporter plasmid containing TetR binding sites upstream of a minimal promoter driving SEAP expression. | Fig. 6<br><br>used in combination with a TetR-Elk1 fusion protein to monitor activity of the MAPK pathway | This work |

**Table S7**: X-ray crystallography data and refinement statistics of 4E1H.95 in complex with 101F Fab

| **4E1H.95/101F Fab** | |
|---|---|
| Wavelength | 1 |
| Resolution range | 49.5 - 2.7 |
| Space group | 65.324 96.212 119.720<br>72 74 70 |
| Unit cell | C 1 2 1 |
| Total reflections | 254,812 (40,624) |
| Unique reflections | 71,869 (11,512) |
| Multiplicity | 3.6 (3.5) |
| Completeness (%) | 99.5 (98.5) |
| Mean I/sigma(I) | 7.7 (1.1) |
| Wilson B-factor ($Å^2$) | 60 |
| R-meas | 0.19 (1.3) |
| CC1/2 | 0.987 (0.398) |
| Reflections used in refinement | 37,066 (3639) |
| Reflections used for R-free | 1854 (182) |
| R-work | 0.216 (0.313) |
| R-free | 0.267 (0.354) |
| Number of non-hydrogen atoms | 7675 |
| Macromolecules | 7646 |
| Solvent | 29 |
| Protein residues | 995 |
| RMS(bonds) | 0.010 |

| | |
|---|---|
| RMS(angles) | 1.59 |
| Ramachandran favored (%) | 95.22 |
| Ramachandran allowed (%) | 4.48 |
| Ramachandran outliers (%) | 0.31 |
| Rotamer outliers (%) | 0.23 |
| Clashscore | 12.6 |
| Average B-factor | 60 |
| macromolecules | 60 |
| solvent | 47 |

**Table S8**: X-ray crystallography data and refinement statistics of 4H.01 in complex with Mota Fab

| 4H.01/Mota Fab | |
|---|---|
| Wavelength | 1.0 |
| Resolution range | 47.1 - 2.8 (2.9 - 2.8) |
| Space group | P $3_2$ 2 1 |
| Unit cell | 90.19 143.56 143.79 120 90 90 |
| Total reflections | 757,244 (117130) |
| Unique reflections | 51,448 (8239) |
| Multiplicity | 14.7 (14.2) |
| Completeness (%) | 99.8 (99.0) |
| Mean I/sigma(I) | 12.6 (1.1) |
| Wilson B-factor (Å$^2$) | 70 |
| R-meas | 0.27 (3.01) |
| CC1/2 | 0.997 (0.56) |
| Reflections used in refinement | 26787 (2633) |
| Reflections used for R-free | 1337 (132) |
| R-work | 0.220 (0.377) |
| R-free | 0.265 (0.394) |
| Number of non-hydrogen atoms | 3928 |
| Macromolecules | 3906 |
| Solvent | 22 |
| Protein residues | 511 |
| RMS(bonds) | 0.013 |
| RMS(angles) | 1.45 |
| Ramachandran favored (%) | 91.25 |
| Ramachandran allowed (%) | 7.55 |
| Ramachandran outliers (%) | 1.19 |
| Rotamer outliers (%) | 0.45 |
| Clashscore | 14.6 |
| Average B-factor | 75 |
| macromolecules | 75 |
| solvent | 60 |

1. D. A. Silva, B. E. Correia, E. Procko, Motif-Driven Design of Protein-Protein Interfaces. *Methods Mol Biol* **1414**, 285-304 (2016).
2. L. Holm, P. Rosenstrom, Dali server: conservation mapping in 3D. *Nucleic Acids Res* **38**, W545-549 (2010).
3. M. B. Keeley, J. Busch, R. Singh, T. Abel, TetR hybrid transcription factors report cell signaling and are inhibited by doxycycline. *Biotechniques* **39**, 529-536 (2005).
